# The role of histone acetyltransferases Gcn5 and Esa1 in recruiting the RSC complex and maintaining nucleosome-depleted regions genome-wide in *Saccharomyces cerevisiae*

**DOI:** 10.1101/2022.06.22.497187

**Authors:** Emily Biernat, Mansi Verma, Matthew Werick, Uzair Khan, Sama Joseph, Chhabi K. Govind

## Abstract

Chromatin remodelers are essential for the maintenance of chromatin structure and gene regulation. In this study, we examined the role of histone acetyltransferases (HATs) Gcn5 and Esa1 in regulating RSC and histone occupancies and their effects on transcription genome-wide. We identified contrasting roles of HATs in modulating RSC occupancies in promoters and ORFs. In HAT mutants, RSC accumulated in nucleosome depleted regions (NDRs) with “fragile nucleosomes (FNs)” more than those with stable -1 nucleosomes. Moreover, the accumulation was more significant in the Esa1 mutant than in the Gcn5 mutant. However, RSC NDR accumulation was not observed in cells lacking H3 or H4 tails. Furthermore, we observed marked increases in histone occupancies in NDRs in the HAT mutants genome-wide. Overall, these data suggest that FNs use hypoacetylated tails to recruit RSC to NDRs, and subsequent acetylation of the tails promote histone eviction. In contrast to the promoters, RSC occupancies were significantly reduced in transcribed ORFs in the HAT mutants. Additionally, the HAT mutants showed reduced TBP and Pol II binding at promoters. Thus, our data implicate HATs and RSC in maintaining NDRs, regulating chromatin structure, and promoting transcription.

## INTRODUCTION

In eukaryotes, DNA is wrapped around a histone octamer (H2A, H2B, H3, and H4) to form the nucleosome. Nucleosomes act as a significant impediment to all stages of transcription (1). One possible mechanism to relieve this impediment is the acetylation of the histone N-terminal tails that weakens the electrostatic interactions between histones and DNA and promotes the recruitment of remodeling complexes that recognize acetylated lysine residues (2, 3). In *S. cerevisiae*, amino-terminal histone tails (NTTs or tails) are acetylated by two major histone acetyltransferase (HAT) complexes: SAGA and NuA4. Gcn5 is the HAT subunit of the SAGA complex and primarily acetylates histone H3. The NuA4 catalytic subunit, Esa1, is the only H4 HAT in the budding yeast and is essential for viability (4–7). Transcription activators such as Gcn4 recruit both SAGA and NuA4 to acetylate nucleosomes and promote histone eviction from promoters (8–11). While it is thought that acetylation promotes transcription, a recent study showed that acetylation in coding regions is a consequence of transcription (12). It is plausible that transcription-dependent acetylation promotes chromatin remodelers’ recruitment or retention to enhance nucleosome accessibility for efficient transcription elongation.

In *Saccharomyces cerevisiae*, the chromatin remodeling complexes SWI/SNF and RSC contain bromodomains that can be utilized to bind acetylated nucleosomes (13–15). These complexes use energy derived from ATP hydrolysis to reposition or evict histones from promoters and stimulate preinitiation complex (PIC) formation (16–18). We have previously shown that RSC is recruited to ORFs of transcriptionally active genes (4, 5) and that RSC-bound nucleosomes are significantly more accessible than non-RSC-bound nucleosomes in transcribed ORFs (16,17,19).

RSC is implicated in maintaining nucleosome depleted regions (NDRs) found upstream of transcription start sites (TSSs) (20–22). RSC evicts nucleosomes formed in NDRs, and slides or evicts nucleosomes flanking the NDRs, termed as -1 and +1 nucleosomes (-1_Nuc and +1_Nuc, henceforth) (21,23,24). As such, the -1_Nuc and +1_Nuc move inward in cells lacking RSC activity, narrowing NDRs (25, 26). Since the TSS for most genes is close to or is part of the DNA wrapped around the +1_Nuc, this nucleosome is either evicted or repositioned during transcription activation to expose the TSS for preinitiation complex (PIC) formation (10,16,27,28). Earlier studies have shown that histones are first hyperacetylated before their eviction during gene activation (27). Considering that promoter nucleosomes are acetylated, the RSC complex could recognize acetylated histones and help in the eviction or repositioning of nucleosomes to promote preinitiation complex assembly (19, 22).

Genome-wide localization studies show that RSC is globally recruited to both -1 and +1 nucleosomes (19,29,30) and to many transcribed ORFs (17,19,24,31). Inactivation of RSC affects multiple stages of transcription, including initiation, TSS selection, elongation, and termination (19,24,26,31–33). Recent studies have also shown that RSC binds to partially-unwrapped nucleosomes found in NDRs known as fragile nucleosome (FNs) (29,34,35). How RSC is recruited to FNs is not known.

The RSC complex contains eight bromodomains (BrDs): Rsc1, Rsc2, and Rsc4 contain two BrDs each; Rsc58 contains one BrD; and one BrD is located at the C-terminus of the ATPase subunit Sth1 (36, 37). RSC BrDs have been shown to bind acetylated lysines (38, 39). For instance, the Sth1 bromodomain binds to acetylated H3K14 *in vitro* and *in vivo* (40–42). Since RSC contains many bromodomains, acetylation may direct RSC recruitment to promoters, including at FNs and in transcribed open reading frames (ORFs) (17,24,43). Recent RSC-nucleosome structures reveal how RSC subunits interact with the nucleosome (44–46). However, the positions of various BrDs remained unresolved. Interestingly, one of the RSC-nucleosome structures revealed the interactions between the H4 tail (unacetylated) with the interface of the ATPase and Snf2 ATP coupling (SnAC) domains. Considering that histone acetylation is shown to prevent octamer transfer of the +1_Nuc by RSC (47), it was suggested that acetylation of the H4 tail might modulate RSC-nucleosome interaction and affect nucleosome sliding or eviction. Thus, both acetylated and unacetylated histone tails could regulate RSC-chromatin interaction and RSC function.

In this study, we determined RSC occupancies in cells lacking either Gcn5, Esa1, or both HATs. We found that RSC accumulated in NDRs genome-wide in cells deficient of HAT function. Further analyses showed that the accumulation is significantly greater in the NDRs that harbor fragile nucleosomes than those with stable -1 nucleosomes (SNs henceforth). We find that RSC accumulation coincides with higher histone occupancies in the same region. Increased RSC occupancies were not seen in cells lacking the H3 or the H4 tail, implying that tails contribute to RSC recruitment/retention in NDRs in cells deficient of HAT functions. In contrast, we saw a significant loss of RSC from transcribed coding sequences in HAT and histone tail mutants. Thus, tails (hypoacetylated or unacetylated) are important for promoting recruitment/retention of RSC to NDRs, especially to FNs, and acetylated histone tails helps to enrich RSC in coding sequences. HAT mutants also show reduced Pol II and TBP occupancies.

## MATERIALS AND METHODS

### Yeast Strain Construction

All *S. cerevisiae* strains used in this study are listed in the supplemental Table S1. Sth1myc-tagged strains were generated as described previously (Govind *et al*. 2012). Sth1 tagged strains were generated using plasmids pFA6a-13xMyc-His3MX6 or pFA6a-13xMyc-TRP1 as the templates to PCR amplify the 13xMyc region using primers containing homologous sequences flanking the stop codon of *STH1,* or primers homologous to the sequences upstream of the region encoding bromodomain of *STH1.* The amplified DNA was used for transformation and colonies were selected for on SC plates deficient for either histidine (SC/His^-^) or tryptophan (SC/Trp^-^).

### Yeast Cell Growth

Cells were grown in YPD at 30°C to an absorbance A_600_ of 0.7-0.8 before being spiked-in and crosslinked. Strains containing the temperature-sensitive *ESA1* allele were grown to an A_600_ of 0.6 before being heat-shocked at 37°C for 1 hour to inactivate Esa1, which were then spiked-in before being crosslinked. Spike-in was performed using *S. pombe*, where *S. pombe* was grown to an A_600_ of 10, and the *S. pombe* culture was added to a final A_600_ of 10% of the A_600_ of the *S. cerevisiae* cultures just before crosslinking. Crosslinking was performed as described previously (Govind *et al.* 2012): 11 ml of crosslinking solution (50mM HEPES-KOH [pH 7.5], 1 mM EDTA, 100 mM NaCl, 11% formaldehyde) was added to 100 ml cultures for a final concentration of 1%. Cultures were then incubated at room temperature for 15 minutes with intermittent shaking before crosslinking was quenched with 15 ml of 2.5M glycine. Cells were collected by centrifugation at 4000 rpm for 5 minutes at 4°C before being washed twice with TBS and stored at -80°C until further use.

### Sonication of Crosslinked Chromatin

Crosslinked cells were thawed on ice and resuspended in pre-chilled 500 μl of FA lysis buffer (50 mM HEPES-KOH [pH 7.5], 1 mM EDTA, 140 mM NaCl, 1% Triton X-100, 0.1% sodium deoxycholate) containing protease inhibitors. Around 500 μl of acid-washed glass beads were added and cells were disrupted for 45 minutes at 4°C. The resulting cell extracts were collected by centrifugation, beads were washed once using 500 μl FA lysis buffer, and the wash was then pooled with the cell extracts. The pooled extracts were then sonicated at 4°C using a Branson Sonifier set at 60% Duty Cycle and an Output Control of 2. Chromatin was sonicated for 12 cycles, where each cycle consisted of active sonication on ice for 30 seconds, followed by a 30-second rest period. The fragmented chromatin was then collected by centrifugation at 4°C for 30 minutes at 15,000 rpm and the supernatant was collected. DNA was purified from 100 μl aliquots of chromatin and checked on a 2% agarose gel to determine the extent of fragmentation. The average size of sonicated DNA was ∼300-400 bp.

### Chromatin Immunoprecipitation

Chromatin immunoprecipitation was performed using a slightly modified protocol (Govind et al. 2012). For myc or Rpb3 ChIPs, 60 μl of anti-mouse Dynabeads (Thermofisher, cat # 11041) were washed twice with PBS/BSA (5 mg/ml BSA) and incubated with 3 μl of anti-Myc (Roche, cat # 11667203001) or anti-Rpb3 (Neoclone, cat # W0012) antibodies in 150 μl of PBS/BSA overnight (anti-Myc) or for 3.5 hours (anti-Rpb3). For H3 or TBP ChIPs, 40 μl of anti-rabbit Dynabeads (Thermofisher, cat # 11-204-D) were washed twice with PBS/BSA and incubated with either 0.5 μl of anti-TBP antibody (gift from Dr. Joseph Reese, Pennsylvania State University) overnight or with 2 μl of anti-H3 (Abcam, cat # ab1791) for 3 hours. The beads were then washed once with PBS/BSA and then twice (once for rabbit-beads) using the following buffers: FA-lysis buffer, wash-buffer II (50mM HEPES-KOH, 500 mM NaCl, 1 mM EDTA, 0.1% sodium deoxycholate, 1% Triton-X 100), and wash-buffer III (10mM Tris-HCl (pH8.0), 250 mM LiCl, 1mM EDTA, 0.5% sodium deoxycholate, 0.5% NP-40 substitute), followed by a final wash with ultrapure water before the elution step. The immunoprecipitated complexes were eluted once using elution buffer I (50mM Tris-HCl, 10mM EDTA, 1% SDS) for 15 minutes at 65°C and then again by using elution buffer II (10mM Tris-HCl, 1mM EDTA, 0.67% SDS) for 10 minutes at 65°C. The two eluents were then combined and incubated overnight in a 65°C hot water bath for reverse crosslinking. Samples were then treated with 5 μl of proteinase K (20mg/ml, Ambion, cat# AM2548) for 2 hours and DNA was extracted twice using chloroform:isoamyl alcohol (IAA). The DNA was then ethanol precipitated overnight at -80°C and resuspended in 50 μl TE/RNase A (10 μg/ml) before being used for library preparation.

### Library Preparation and Sequencing

ChIP DNA (7-10 ng) was processed for library preparation using the NEBNext Ultra II Library prep kit for Illumina (cat # E7465S) to generate ChIP-seq libraries. NEXTFLEX ChIP-Seq barcode oligos from BIOO Scientific (cat# NOVA-514122) were used for ligation and the libraries were amplified according to the manufacturer instructions, where the ChIP samples were amplified for 9-15 cycles. The DNA libraries were gel purified and sequenced on the Illumina HiSeq platform in paired-end mode at GENEWIZ in South Plainfield, NJ, USA.

### Data Analysis

Sequences were trimmed to remove adapters using Cutadapt (parameter: -m 20) and were aligned to the *S. cerevisiae* (SacCer3) genome using Bowtie2 (parameters: -X 1000 –very-sensitive –no-mixed –no-unal). Samtools was used to sort, index, and remove PCR duplicates from the .bam files. The ChIP-seq .bam files were analyzed using BamR (https://github.com/rchereji/bamR). The ChIP-seq reads were normalized to a value of 1 for each chromosome. Metagene profiles and scatterplots were generated using JMP and boxplots were generated using the website http://shiny.chemgrid.org/boxplotr/. The list of fragile and stable nucleosomes was taken from a previous study (25). The average occupancies in ORFs, at NDRs, and at -1 and +1 nucleosome positions were calculated using BamR scripts.

## RESULTS

### RSC accumulates in promoters in the HAT mutants

To determine whether acetylation by SAGA or NuA4 promotes the recruitment of RSC to chromatin (Fig. 1A), we performed Sth1-Myc (RSC) ChIP-seq in wild type (WT) *S. cerevisiae* cells and in the *sth1ΔBrD*, *gcn5Δ*, *esa1ts* (esa1 hereafter) and *esa1/gcn5Δ* mutants. The *esa1* and *esa1/gcn5Δ* cells were heat-shocked (HS) to inactivate Esa1. *gcn5Δ* and *esa1* showed significantly reduced H3 and H4 acetylation (Fig. S1A and S1B), as shown previously (8, 10).

**Figure 1:**
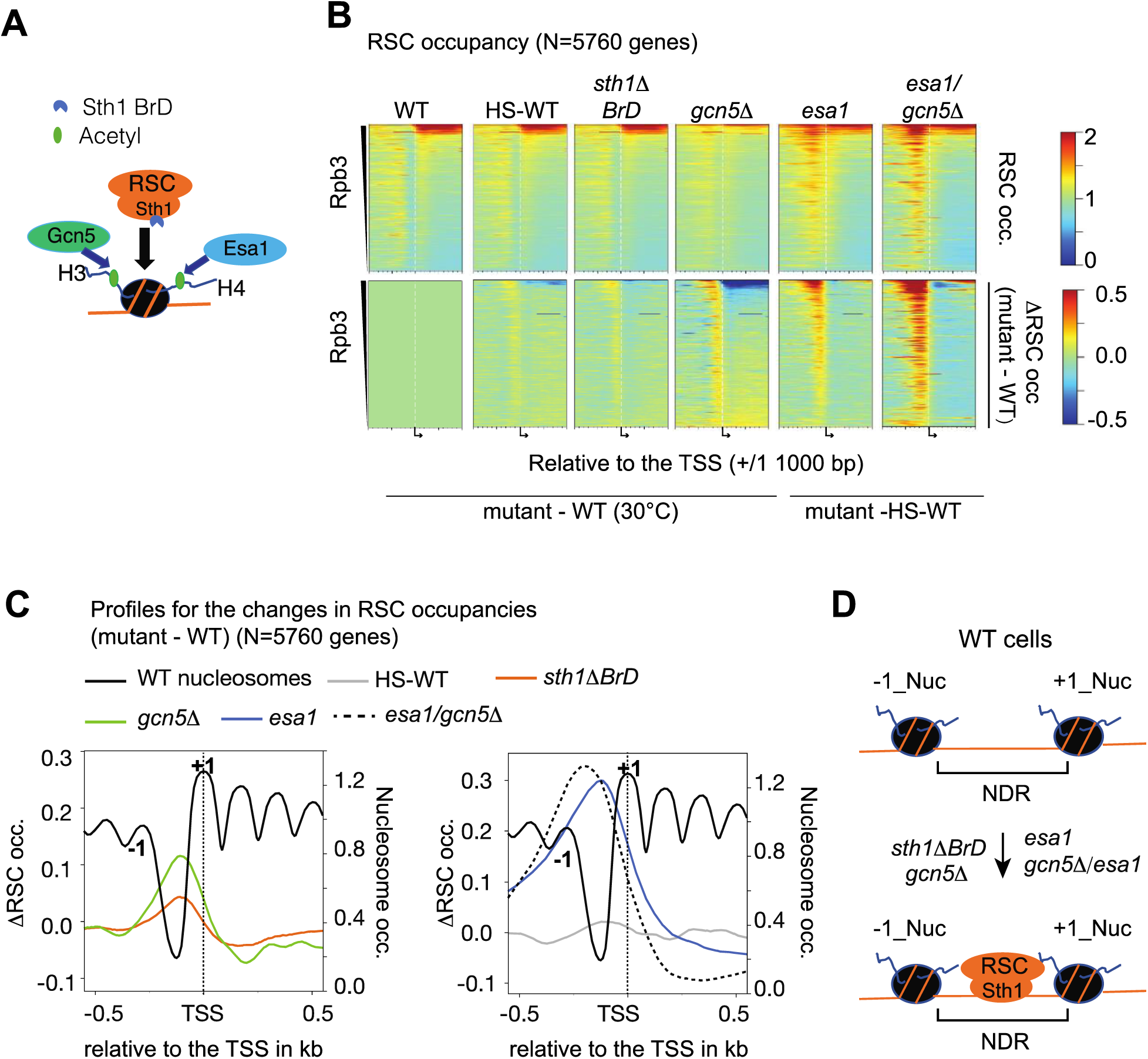
RSC occupancy increases in NDRs in cells lacking HATs or the Sth1 bromodomain. A) Schematic showing that acetylation of histone tails promotes recruitment of RSC to chromatin (nucleosomes). The H3 tail is acetylated by Gcn5 and the H4 tail by Esa1. B) Heatmaps in the top panels show RSC occupancies (RSC occ.) and bottom panels show changes in RSC occupancies (mutant – WT or mutant – HS-WT; ΔRSC occ.) at 5760 genes for WT cells (30°C), heat-shocked WT cells (HS-WT), *gcn5Δ*, *sth1ΔBrD* (lacks C-terminal 1270-1359 residues), *esa1* (*esa1-L254P; esa1ts*) and *esa1/gcn5Δ* cells. Genes were sorted by decreasing average Pol II (Rpb3) occupancies in their ORFs. The heatmaps show +1/-1000 bp of the transcription start sites (TSS). C) Metagene profiles showing RSC occupancy changes at all genes (n=5760) in indicated strains. The left-hand y-axis shows ΔRSC occ. and the right-hand y-axis shows the nucleosome profile of WT cells (data taken from (19)). Data for +/-500 bp of TSS is shown. +1 and -1 represent the -1 and +1 nucleosome positions. D) Schematic showing that in RSC occupancies increase in the NDRs of cells lacking HATs (Gcn5, Esa1, or both) or Sth1 BrD. The -1 and +1 nucleosomes are represented as -1_Nuc and +1_Nuc, respectively.

We plotted RSC occupancies (Fig. 1B, top panels) and changes in RSC occupancies (mutant – WT) (Fig. 1B; bottom panels) as heatmaps. The WT cells showed RSC enrichment in promoters, including at -1_Nuc and +1_Nuc positions, consistent with previous studies (Fig. 1B; top left panel, yellow hues near the TSS) (23,29,48). In addition, RSC was enriched in the coding regions of highly transcribed genes (Fig. 1B, top of the WT heatmap), as shown previously (17,19,24,31). RSC has been shown to redistribute from ORFs to promoters when cells are subjected to heat shock (31). We also saw a slight increase in RSC occupancy upstream of the TSS in heat-shocked WT cells (HS-WT) and loss from the ORFs (Fig. 1A; compare the top and the bottom panels for WT and HS-WT). Thus, expected RSC occupancies were seen for both WT and HS-WT cells.

Since RSC BrDs bind acetylated histone tails (15, 40), we expected to see lower RSC occupancies in cells lacking Gcn5 or with diminished Esa1 function. Instead, RSC occupancies increased in the promoters across the genome in the mutants to varying extents (Figure 1B-C). *sth1ΔBrD* showed a slight enrichment upstream of the TSS, resembling the increase in HS-WT cells (Fig. 1B-C). The increases were more significant in the *gcn5Δ* mutant (Fig. 1B; compare yellow/red hues upstream of TSS in WT, *sth1ΔBrD* and *gcn5Δ*, and Fig. 1C). Strikingly, the RSC occupancies were substantially higher in the *esa1* and *gcn5Δ/esa1* mutants compared to other mutants (Fig. 1B and 1C). The RSC accumulation seen in these mutants is unlikely to be an effect of heat shock because the changes in RSC occupancies for the *esa1* and *esa1/gcn5Δ* mutants were calculated against HS-WT (Fig. 1C).

Plotting RSC occupancy changes (ΔRSC: mutant -WT) along with the MNase-seq nucleosome occupancy profile for WT cells revealed that RSC was predominantly accumulating in NDRs in the mutants (Fig. 1C). The profiles reveal that RSC also increases over the -1_Nuc in the *esa1/gcn5Δ* double mutant. Contrary to the expectation that RSC occupancies would decrease in the mutants, our data show increases in NDRs, suggesting that Gcn5 and Esa1 prevent accumulation of RSC in promoters and in NDRs, genome-wide (Fig. 1D).

### RSC accumulates in NDRs containing fragile nucleosomes in HAT deficient cells

We next asked whether the RSC occupancy increases correlate with the width of NDRs. We did not see any significant correlation between the NDR widths of genes and RSC promoter occupancies in HS-WT, *sth1ΔBrD* and *gcn5Δ* cells (Fig. 2A). However, we did observe slightly higher correlations in the *esa1* and *esa1/gcn5Δ* mutants, suggesting that wider NDRs might be prone to RSC accumulation in these mutants.

**Figure 2:**
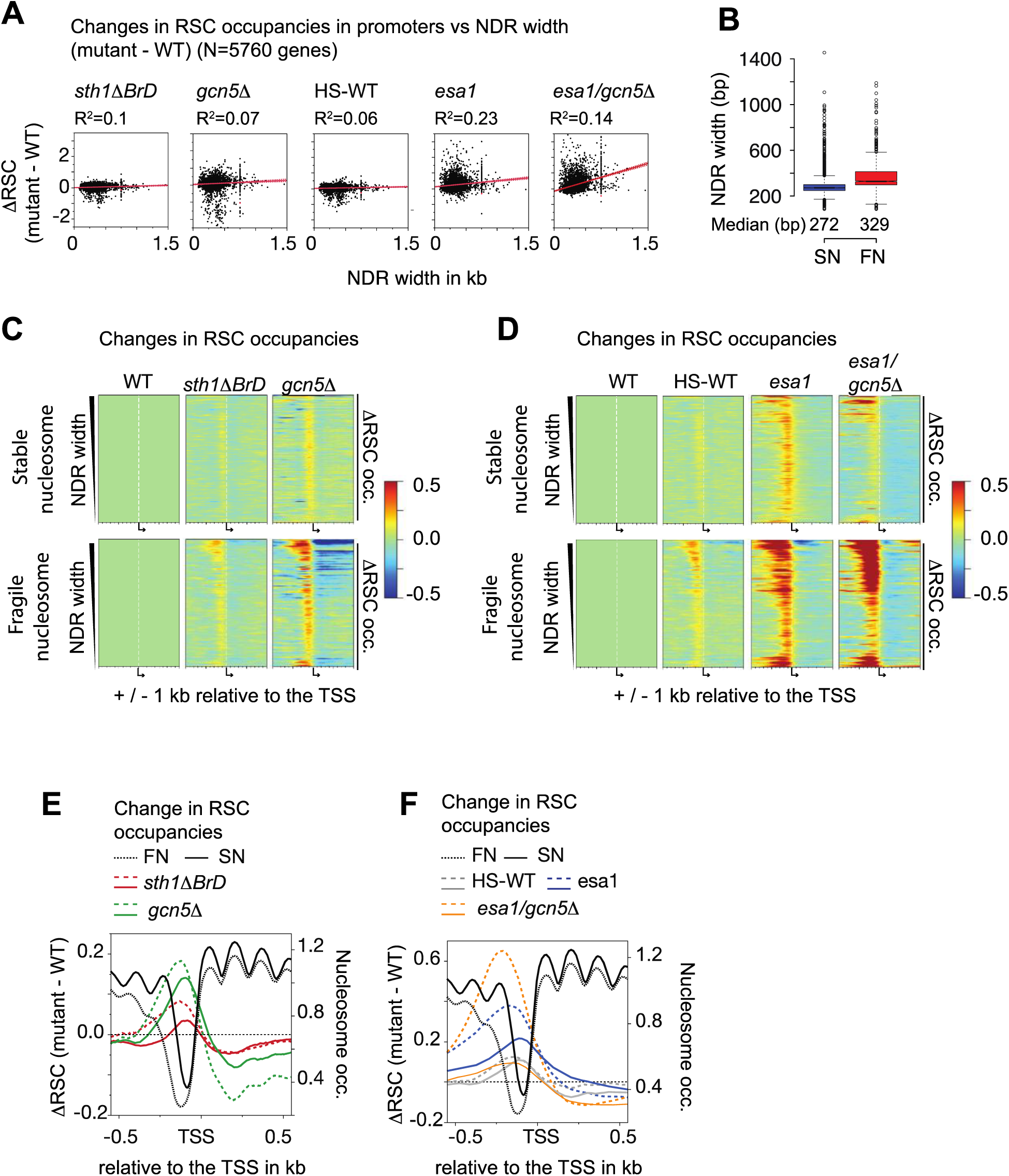
RSC accumulates in the promoters of genes containing fragile nucleosome in cells lacking HATs. A) Scatterplots showing the correlations between NDR width and the changes in RSC occupancies in promoters. The RSC occupancies were calculated for 350 bp upstream of the TSS for each gene. Scatterplots are shown for *sth1ΔBrD, gcn5Δ, gcn5Δ/sth1ΔBrD, esa1,* and *gcn5Δ/esa1*. B) Boxplot depicting the NDR widths for genes in which the NDRs contain fragile nucleosomes (FNs; N=1888) along with genes containing stable nucleosomes within their NDRs (SNs; N=2975). Median length in bp is shown for FN-NDRs and SN-NDRs. The list for FNs and SNs was taken from (25, 49). C-D) Heatmaps showing the changes in RSC occupancies (ΔRSC occ., mutant – WT) at NDRs containing stable nucleosomes (top panels) or fragile nucleosomes (bottom panels) in *sth1ΔBrD* and *gcn5Δ* cells (C), and HS-WT, *esa1* and *esa1/gcn5Δ* cells (D). The control heatmap (WT-WT) is shown on the left for each figure. Genes were sorted by decreasing NDR widths and centered at the TSS. E-F) Metagene profiles showing the changes in RSC occupancies in WT, *sth1ΔBrD* and *gcn5Δ* (F) and HS-WT, *esa1*, *esa1/gcn5Δ* cells (G). Solid traces (–) represent occupancy changes for SN-NDR genes, and dashed (--) traces show changes for the FN-NDRs. Horizontal dotted traces show no change. The left-hand y-axis shows ΔRSC, and the right-hand y-axis shows nucleosome occupancies calculated using MNase-seq (19).

RSC has been shown to bind to partially-unwrapped FNs (25, 35), which are generally found in wider NDRs (25). As such, NDRs shown to harbor FNs were significantly wider than NDRs that harbor stably bound SNs at the -1 position upstream of the TSS (Fig. 2B). To test the idea that RSC might be accumulating in NDRs containing FNs, we separated genes based on the presence of FNs or SNs within their promoters (25, 49). There were 1887 genes with -1_FN-containing NDRs (FN-NDRs hereafter) and 2974 genes with -1_SN-containing NDRs (SN-NDRs hereafter).

The heatmaps showing ΔRSC at FN-NDRs and SN-NDRs revealed a general global pattern: RSC accumulation was significantly greater in FN-NDRs than SN-NDRs (Fig. 2C and 2D; compare the top panel of each mutant with the corresponding bottom panel). The ΔRSC profiles show that the peak changes were centered within the NDRs for both SNs and FNs, reinforcing the idea that RSC accumulates in NDRs in *sth1ΔBrD* and *gcn5Δ* cells (Fig. 2E). The differences between the accumulation of RSC in SN-NDRs vs. FN-NDRs were striking in the e*sa1* and *esa1/gcn5Δ* mutants compared to the *gcn5Δ* or *sth1ΔBrD* mutants (Fig. 2C and 2D; compare top panels with corresponding bottom panels). RSC accumulated to a greater extent in FN-NDRs than in SN-NDRs in the *esa1* mutant (Fig. 2F; compare solid blue to dashed blue trances). This suggests that in cells lacking the H4 HAT Esa1, RSC is more prone to accumulation in NDRs with fragile nucleosomes than in those with stable nucleosomes.

The *esa1/gcn5Δ* double mutant showed the most significant differences in RSC occupancies between SN-NDRs and FN-NDRs (Fig. 2F; compare solid and dashed orange traces). The ΔRSC profiles show that the RSC increase at SN-NDRs was smaller in *esa1/gcn5Δ* than in *esa1*, and was comparable to the accumulation seen in the HS-WT cells (Fig. 2F; compare solid gray, orange, and blue traces). In contrast, RSC accumulates considerably at FN-NDRs in *esa1/gcn5*Δ, much more than that seen in either of the *esa1* or *gcn5*Δ single mutants (compare dashed green traces in Fig. 2E with the dashed blue and orange traces in Fig. 2F). Since nucleosomes are hypoacetylated in HAT mutants compared to WT cells (Fig. S1), our data suggest that hypoacetylated/unacetylated fragile nucleosomes recruit/retain RSC more efficiently than hypoacetylated or unacetylated stable nucleosomes. Moreover, the greater RSC accumulation in FN-NDRs in *esa1* cells compared to *gcn5Δ* cells suggest that H4-hypoacetylated FNs are better at retaining/recruiting RSC than H3-hypoacetylated FNs.

### Sth1 bromodomain and HATs Gcn5 and Esa1 stimulate histone eviction from NDRs

We next determined histone H3 occupancies by ChIP-seq in the HAT and Sth1 BrD mutants to ask whether RSC accumulates on histones in promoters. We calculated the average H3 occupancies for each gene at the region encompassing -1_Nuc, NDR, and +1_Nuc positions (-1,NDR,+1 hereafter). We found that the average H3 occupancies increased significantly at -1,NDR,+1 in *sth1ΔBrD* and *gcn5Δ* relative to WT, and in *esa1* and *esa1/gcn5Δ* cells relative to HS-WT (Fig. S2A). The heatmaps depicting the changes in H3 occupancies (mutant – WT; ΔH3 occ.) show that H3 globally increased in NDRs to varying degrees in different mutants (Fig. 3A, compare the yellow hues upstream of the TSSs in the mutants). Compared to a relatively small increase in HS-WT, the H3 increases were significantly greater in both the *sth1ΔBrD* and *gcn5*Δ mutants. In the *gcn5*Δ mutant, histone occupancies increased slightly at -1 and +1 positions at these genes (Fig. 3A, top of heatmap). This was not evident in *sth1ΔBrD*, suggesting that Gcn5 promotes the eviction of the +1 and -1 nucleosomes of highly transcribed genes, and of nucleosomes from NDRs globally (Figs. 3A and 3B).

**Figure 3:**
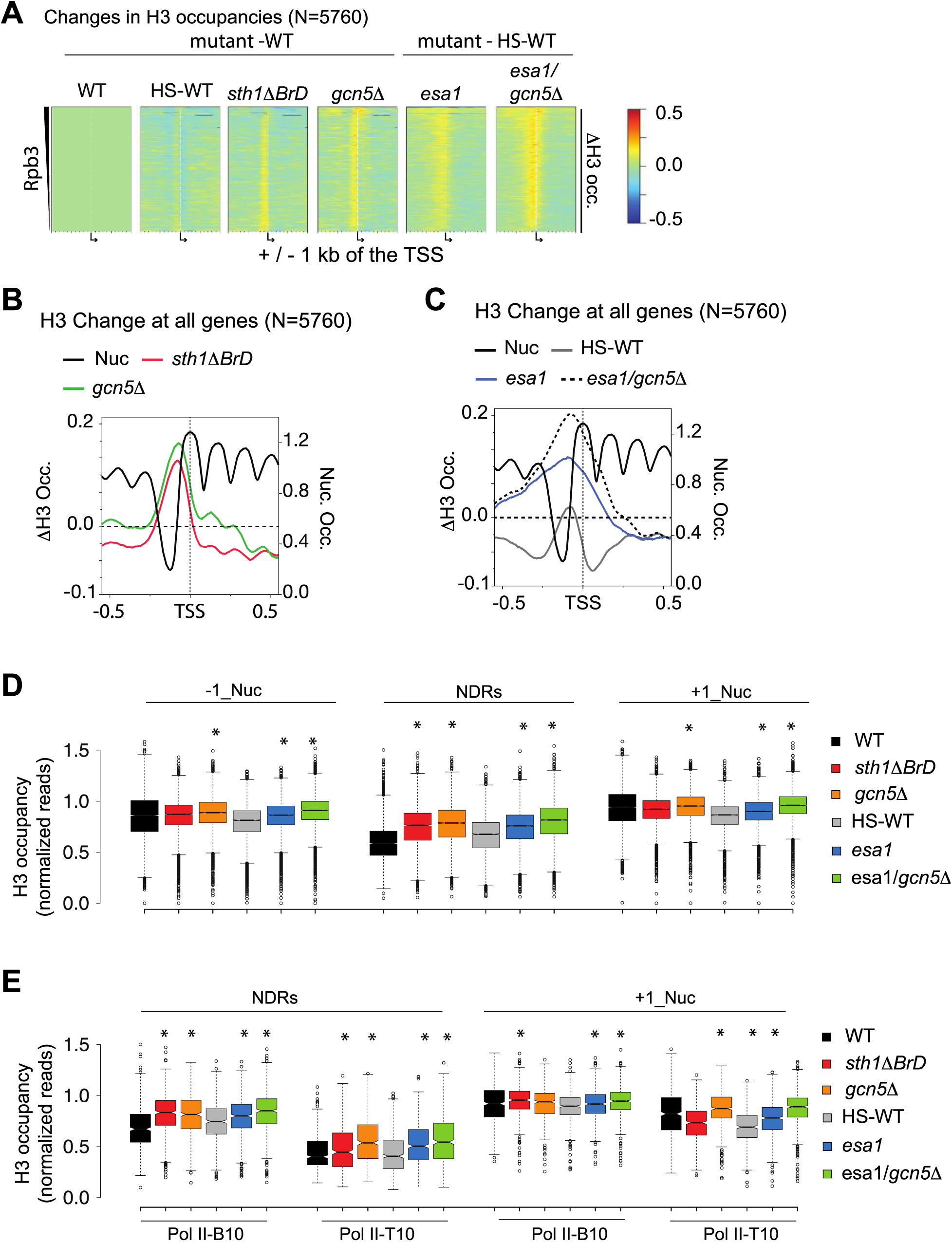
Loss of HATs Gcn5 and Esa1 increase H3 occupancies in NDRs. A) Heatmaps showing changes in H3 occupancies (mutant – WT, Δ H3 occ.) are shown for *sth1ΔBrD*, *gcn5Δ*, HS-WT, *esa1* and *esa1/gcn5Δ* cells. The control heatmap (WT-WT) is shown on the left. Genes are sorted in decreasing order of average Pol II occupancies in their ORFs and are centered at the TSS. For *esa1*-containing mutants, the changes were calculated using heat shocked WT (HS-WT) H3 ChIP-seq data. B-C) Metagene profiles showing the changes in H3 occupancies (ΔH3 occ.) at 5760 genes for *sth1ΔBrD* and *gcn5Δ* (B), and for HS-WT, *esa1* and *esa1/gcn5Δ* (C). The changes are plotted +/- 500 bp around the TSS. The ΔH3 occ. are plotted on the left-hand y-axis, and nucleosome occupancies (Nuc occ.) are plotted on the right-hand y-axis. D) Boxplot comparing the H3 occupancies for all genes (N=5760) at the -1 nucleosome (-1_Nuc), NDR, and +1 nucleosome (+1_Nuc) positions in WT, *sth1ΔBrD*, *gcn5Δ,* HS-WT, *esa1* and *esa1/gcn5Δ*. Asterix (*) denotes significant changes (p-value < 0.05) in the mutant relative to the WT (or HS-WT for *esa1*-containing mutant). The Welch t-test was performed for p-values. E) Boxplot showing H3 occupancies for the top and bottom 10% of transcribed genes (Pol II-T10 and Pol II-B10, respectively) at NDRs and the +1_Nuc position in WT, HS-WT, *sth1ΔBrD*, *gcn5Δ, esa1* and *esa1/gcn5Δ* cells. Significant H3 increases compared to WT cells were determined using the Welch t-test and are denoted by an asterisk (*).

Likewise, *esa1* and *esa1/gcn5Δ* showed higher H3 occupancies in NDRs (Figs. 3A and 3C). However, unlike those seen in the *gcn5*Δ and *sth1*Δ*BrD* mutants, these increases were not restricted to NDRs, but instead extended farther upstream and downstream of the NDRs (Fig. 3A, the two panels in the right-hand side, and Fig. 3C). Thus, the data suggest that both HATs and Sth1 BrD help evict nucleosomes that might assemble within NDRs.

We next inspected the changes in global H3 occupancies at the -1_Nuc position, +1_Nuc position, and at NDRs individually for each mutant. All mutants showed substantially higher H3 occupancies in NDRs, and at -1_Nuc and +1_Nuc positions except *sth1ΔBrD* (Fig. 3D). Interestingly, both *sth1ΔBrD* and *gcn5Δ* mutants showed very similar increases in H3 occupancies at NDRs. Thus, Sth1 bromodomain and HATs are important for keeping NDRs depleted of histones, genome-wide.

Activators recruit SAGA and NuA4 to promote histone eviction and to stimulate transcription of their target genes (10,16,50–52). Eviction of histones from promoters and repositioning of +1 nucleosome exposes the TBP binding site (26). As such, the HAT mutants might show more significant increases in H3 occupancies at highly transcribed genes than at lowly transcribed genes. Therefore, we compared the H3 occupancies at the top 10 (Pol II-T10) and the bottom 10 (Pol II-B10) percent transcribed genes (N=539) at NDRs and at the +1_Nuc position (Fig. 3E). At the NDRs of Pol II-B10 genes, H3 occupancy in the *sth1ΔBrD* mutant was comparable to or even slightly higher than in *gcn5Δ* (Fig. 3E; compare Pol II-B10 NDRs data for *sth1ΔBrD* and *gcn5Δ*). Thus, Sth1 BrD acts on par with Gcn5 in evicting nucleosomes from the NDRs of the least transcribed genes. At Pol II-T10 NDRs, H3 occupancies were higher in *gcn5Δ*, *esa1*, and *esa1/gcn5Δ* cells compared the *sth1ΔBrD* mutant (Fig. 3E, compare H3 at Pol II-T10 genes in WT, *sth1ΔBrD* and *gcn5Δ*, and HS-WT, *esa1* and *esa1/gcn5*). Thus, Gcn5 and Esa1 make greater contributions in evicting nucleosomes than Sth1 BrD, and Sth1 BrD plays only a minor role in evicting histones from NDRs of highly expressed genes. However, we find that the deletion of Sth1 bromodomain increases H3 occupancies at -1,NDR,+1 for all but the top 25% of transcribed genes (Fig. S2B), implying that the Sth1 bromodomain contributes to histone eviction from promoters at the majority of genes.

At Pol II-B10 genes, the increase in H3 occupancy at the +1_Nuc position were less pronounced than those seen in NDRs (Fig. 3E, compare NDRs vs. Pol II-B10 +1_Nuc). Only *sth1ΔBrD*, *esa1*, and *esa1/gcn5Δ* showed significant H3 increases at the +1_Nuc location for lowly transcribed genes. At the Pol II-T10 genes at +1_Nuc position, H3 occupancies were higher in *gcn5Δ* than *esa1*, and the highest occupancy was seen in the *esa1/gcn5Δ* double mutant (Fig 3E; right-hand panel). Thus, the data suggest that Sth1 BrD plays a greater role in evicting histones from NDRs of less transcribed genes, while the HATs are more important for evicting histones from the NDRs and +1 position of highly transcribed genes (Fig. 3E). In fact, *gcn5Δ* and *esa1/gcn5Δ* showed significantly greater increases in H3 occupancies at promoters of RPGs than the top 20% of transcribed genes excluding RPGs (Fig. S2C and S2D). Thus, HATs help in clearing nucleosomes from highly transcribed genes, potentially allowing them to achieve higher expression levels.

### HAT mutants evoke comparable increases in histone occupancies in NDRs containing stable vs. fragile nucleosomes

Since FNs-NDRs were more enriched for RSC than SN-NDRs in the mutants (Fig 2C-2F), we next examined histone occupancies at -1,NDR,+1 at SNs and FNs. H3 occupancies were higher in SN-NDRs compared to FN-NDRs (Fig. 4A; compare stable and fragile nucleosomes, black boxes). Both *sth1ΔBrD* and *gcn5Δ* showed higher H3 occupancies than WT at SN-NDRs and FN-NDRs (Fig. 4A), however, the increases were higher in the *gcn5Δ* compared to *sth1ΔBrD* cells in both types of NDRs. Although both *esa1* and *esa1/gcn5Δ* showed higher H3 occupancies than the HS-WT cells, the highest increase was seen in *esa1/gcn5Δ*. The greater histone occupancies in *esa1/gcn5Δ* suggest that both Gcn5 and Esa1 aid in efficient histone eviction from promoters.

**Figure 4:**
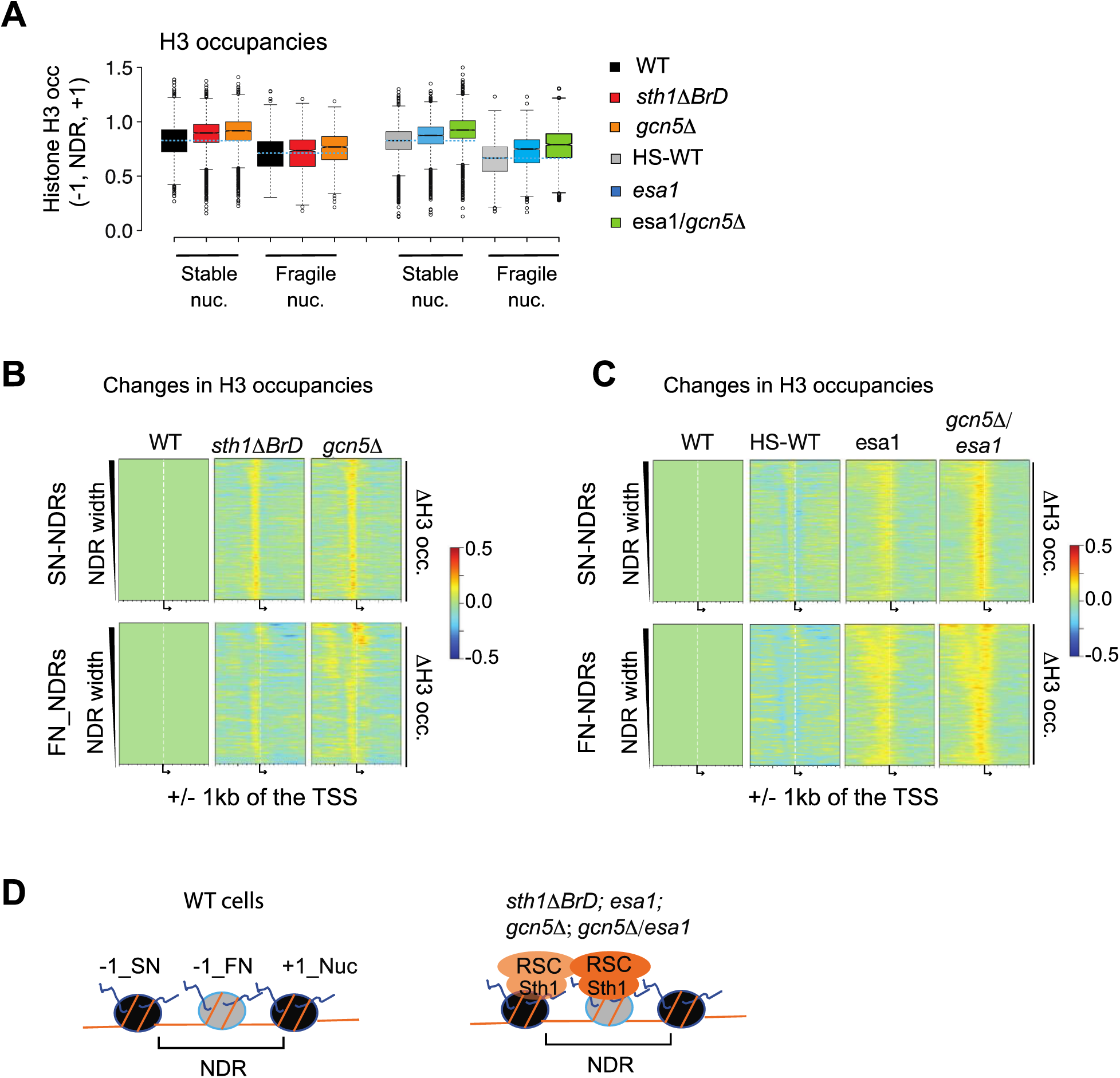
Histone occupancies increase differently in NDRs containing stable vs. fragile nucleosomes. A) Boxplot showing the normalized H3 occupancies at promoters (-1_Nuc, NDR, and +1_Nuc positions) of FNs and SNs in WT cells, HS-WT cells, *sth1ΔBrD* and *gcn5Δ*, *esa1* and *esa1/gcn5Δ* cells. The dotted blue line indicates the median of either WT or HS-WT H3 occupancies for comparison. B-C) Heatmaps showing the changes in H3 occupancies (mutant – WT) at NDRs containing stable nucleosomes (top panels) and fragile nucleosomes (bottom panels) in *sth1ΔBrD* and *gcn5Δ* cells (B), and in HS-WT, *esa1* and *esa1/gcn5Δ* cells (C). The control heatmap (WT-WT) is shown on the left. Genes were sorted by decreasing NDR widths and centered at the TSS. D) A schematic showing that histone accumulation in NDRs also traps RSC on fragile (-1_FN) and stable (-1_SN) nucleosomes present at -1 positions. The lighter RSC color (orange) shade at -1_SN represents lower RSC occupancy than at -1_FN.

The ΔH3 heatmaps at SN-NDRs were very similar to ΔRSC (compare heatmaps in Figs. 4B and 4C top panels with Figs. 2C and 2D top panels). Compared to SN-NDRs, the H3 increases were slightly less pronounced at FN-NDRs in the *sth1ΔBrD* and *gcn5Δ* mutants (Fig. 4B, compare the top panels with the corresponding bottom panels). The *esa1* and *esa1/gcn5Δ* to show a slightly greater increase in H3 for FN-NDRs than SN-NDRs (Fig. 4C).

At SN-NDRs, the higher RSC and H3 occupancies in the HAT and Sth1 BrD mutants (the top panels of Fig. 2C and 2D, and 4B and 4C) might suggest that there is more RSC in NDRs simply because there are more histones for RSC to interact with. However, in FN-NDRs of the *Sth1ΔBrD* and HAT mutants, the RSC increases were much higher than the H3 increases (compare bottom panels of Fig 2C and 2D with bottom panels of 4B and 4C, respectively). The relatively higher RSC occupancies at FN-NDRs than H3 might suggest that fragile nucleosomes recruit or retain more RSC than stable nucleosomes (Fig. 4D). Since FN-NDRs display lower histone occupancies than SN-NDRs promoters (Fig. 4A, compare WT H3 at SN and FN-NDRs), it is possible that the ability of FNs to recruit RSC more efficiently helps in maintaining lower histone occupancies at FN-NDRs than SN-NDRs.

### RSC accumulation in promoters/NDRs is histone tail-dependent

Considering that Gcn5 and Esa1 acetylate histone tails and that RSC accumulates in NDRs of cells lacking Esa1 and Gcn5, we asked whether the accumulation of RSC is histone tail-dependent. To test this, we determined RSC occupancies in the histone tail deletion mutants *H3Δ1-28* and *H4Δ1-16*. The deleted regions contain several acetyl-lysines: K9, K14, K18, K23, and K27 for the H3 tail and K5, K8, K12, and K16 for the H4 tail. Since the HAT mutants show higher RSC occupancies in NDRs, we expected the tail mutants to also display increased RSC occupancies. However, the RSC occupancies appear to decrease in *H3Δ1-28*, and only slightly increase in *H4Δ1-16* (Fig. 5A)

**Figure 5:**
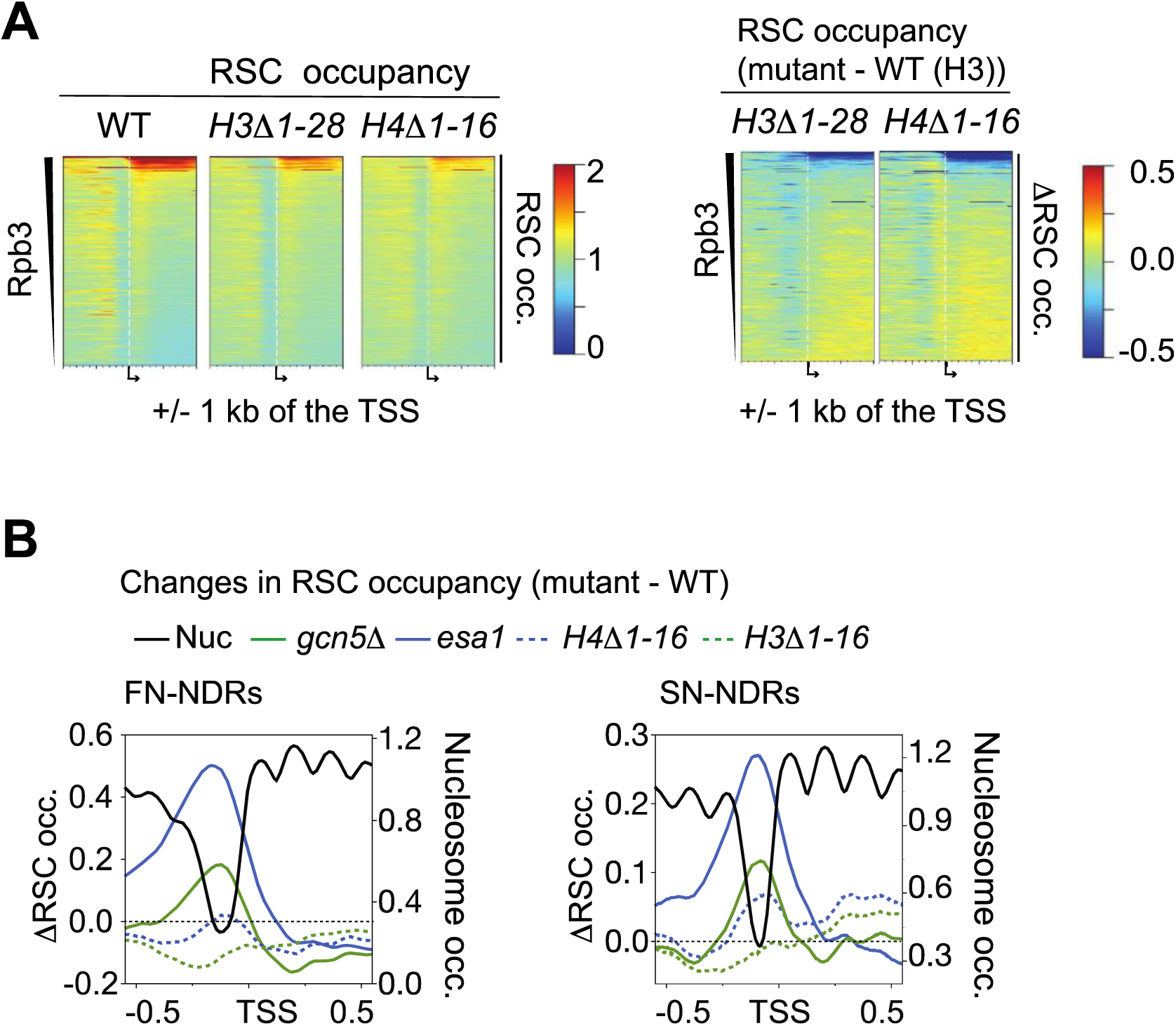
RSC does not increase in NDRs in cells lacking H3 or H4 N-terminal tails. A) Heatmaps depicting RSC occupancies for all genes in WT cells and in cells lacking either the H3 or H4 N-terminal tail (NTT) (first three heatmaps), and RSC occupancy changes (ΔRSC occ., mutant – WT, last two heatmaps on the right hand). Genes are sorted by decreasing average Pol II occupancies in their ORFs and are centered at the TSS (N=5760). B) Metagene profiles comparing ΔRSC occupancies (ΔRSC occ.) in histone NTT mutants with *esa1* and *gcn5Δ* mutants at FN-NDRs (left-hand panel) and SN-NDRs (right-hand panels).

We next compared the differences between the ΔRSC profiles of the *H3Δ1-28* and *H4Δ1-16* mutants vs. ΔRSC profiles of the HAT mutants at SN and FN NDRs. Comparing ΔRSC in *gcn5Δ* vs. *H3Δ1-28* (Fig. 5B; compare solid and dotted green traces) and in *esa1* vs. *H4Δ1-28* (compare solid and dotted blue traces), it is clear that the increases seen in the HAT mutants were largely absent in the tail mutants. Moreover, the differences were larger between the tail mutants and the HAT mutants at FN-NDRs than SN-NDRs (Fig. 5B; compare solid traces with the dotted traces in both panels; note the changes in y-axis scales). The ΔRSC in *esa1* vs. *H4Δ1-16* were larger than ΔRSC in *gcn5Δ* vs. *H3Δ1-28* mutants (Fig. 5D; compare the differences between solid and dotted traces).

The observation that RSC occupancies increase in *gcn5Δ* and *esa1* mutants, but not in the tail mutants, suggests a role for unacetylated or hypoacetylated tails in this process. A recent RSC-nucleosome Cryo-EM structure revealed that the (unacetylated) H4 tail binds to the interface of the SnAC and ATPase domains of Sth1 (46). The binding of the tail could stabilize the RSC-nucleosome interaction, and this enhancement of the RSC-nucleosome interaction by the unacetylated tail would explain why RSC accumulates when HATs are impaired and not in WT or *H4Δ1-16*. In WT cells, acetylation of the H4 tail might weaken the RSC-nucleosome interaction and could provide a potential mechanism for disengaging RSC from chromatin. Since *H3Δ1-28* did not show accumulation of RSC as seen in *gcn5Δ* (Fig. 5B), we speculate that the H3 tail might also contribute to the retention of RSC in NDRs. We propose that unacetylated/hypoacetylated nucleosomes bring more RSC to FN-NDRs, promoting efficient nucleosome eviction and maintaining nucleosome-depleted regions.

### HATs Gcn5 and Esa1 promote RSC Recruitment to transcribed ORFs

We and others have previously shown that RSC is enriched in many transcribed ORFs, in addition to promoters (16,17,19,24,31). We analyzed the effect of HAT mutants on RSC occupancies in ORFs. In agreement with our previous studies, RSC ORF occupancies strongly correlated with that of Pol II (Rpb3) occupancies, genome-wide (R^2^= 0.78; Spearman ρ = 0.78) (Fig. S3A), supporting the idea that RSC is co-transcriptionally recruited to coding regions. The correlations were significantly reduced in both *gcn5*Δ (R^2^= 0.49; Spearman ρ = 0.62) and *esa1* (R^2^= 0.48; Spearman ρ = 0.66), but not in *sth1ΔBrD* (R^2^= 0.79; Spearman ρ = 0.78) (Fig. S3A).

Considering that RSC is recruited to highly transcribed ORFs, we analyzed the top 20% Pol II-occupied genes (Pol II-T20; N=1092) in greater detail (Fig. 6A and 6B). Contrary to the increased RSC occupancies in NDRs, the RSC occupancies in ORFs were substantially reduced in the *gcn5*Δ mutant (Fig. 6A; top right of the heatmap, Fig. S3B; compare *gcn5Δ* heatmap with WT heatmap, S3C), indicating that Gcn5 is needed for recruiting or retaining high RSC occupancies over the ORFs of transcribed genes. Although acetylated H3K14 (acetylated by Gcn5) enhances the binding of the Sth1 BrD to nucleosomes *in vitro* and *in vivo* (40–42), *sth1ΔBrD* showed minimal reductions in RSC occupancy compared to *gcn5Δ*. The minor reductions in RSC occupancies in the *sth1ΔBrD* mutant suggest that other RSC bromodomains might act redundantly to recruit RSC to transcribed ORFs. Significant reductions in occupancies were also seen in *esa1* and *gcn5Δ/esa1* mutants (Fig. 6B). However, reductions were modest compared to *gcn5Δ*. RSC recruitment defects in *esa1* and *esa1/gcn5Δ* mutants might be dampened by increased interactions between the H4 tail and RSC, as observed at NDRs (Figs. 1-5). Nonetheless, our data show that Gcn5 promotes RSC enrichment in transcribed ORFs, and Esa1 also contributes to this process.

**Figure 6:**
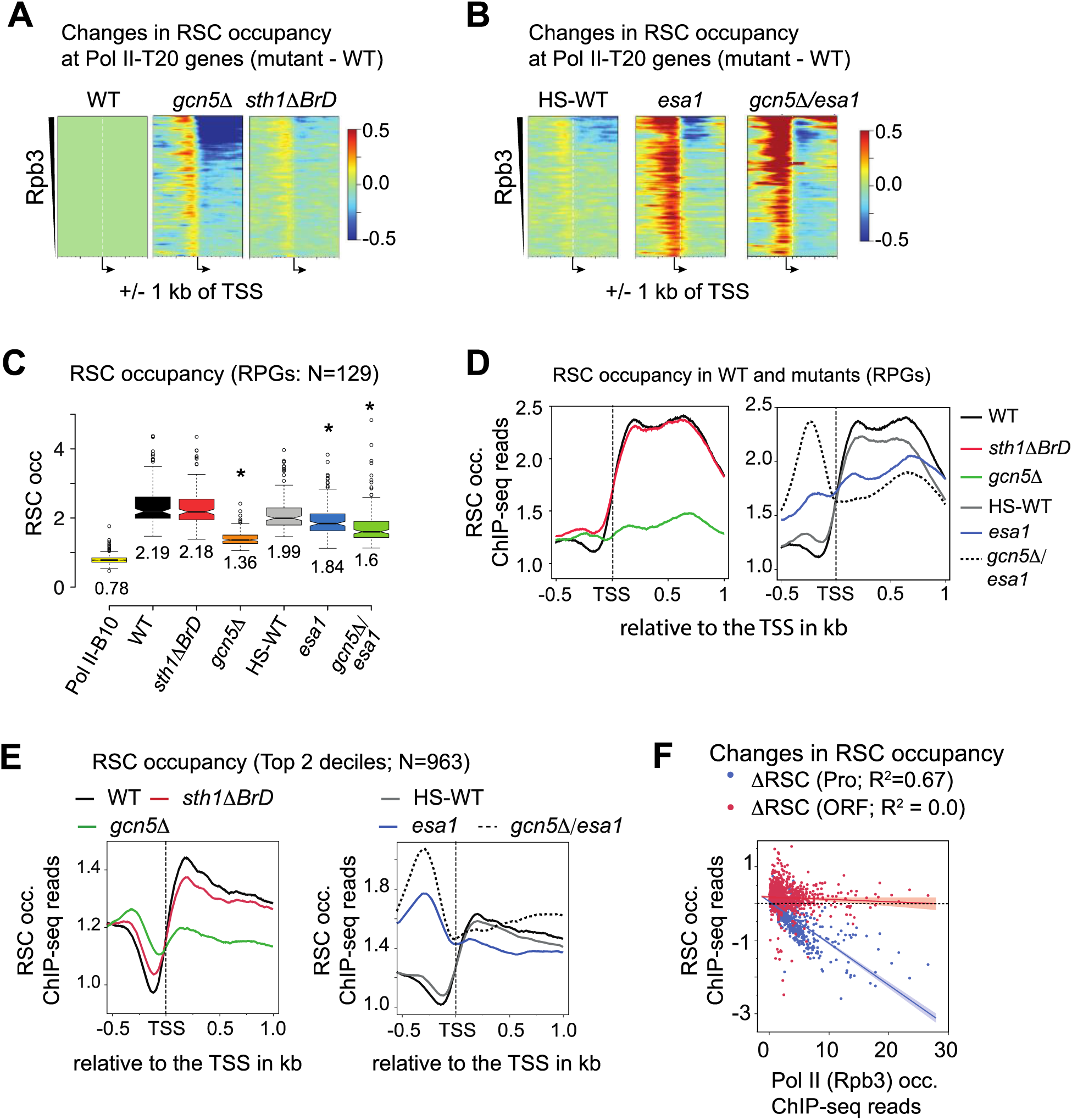
HATs Gcn5 and Esa1 promote RSC enrichment in transcribed ORFs. A-B) Heatmaps showing the changes in RSC occupancies (mutant – WT) at the top 20% Pol II-occupied genes (Pol II-T20). The changes are shown for the *sth1ΔBrD* and *gcn5Δ* cells (A), and for HS-WT, *esa1* and *esa1/gcn5Δ* cells (B). The control heatmap (WT-WT) is shown on the left in (A). The genes were sorted by decreasing average Pol II occupancies in their ORFs. The heatmaps show +/- 1000 bp of the transcription start sites (TSS). C) Boxplot depicting the normalized RSC occupancies in WT, HS-WT, *sth1ΔBrD*, *gcn5Δ*, *esa1* and *esa1/gcn5Δ* cells at Ribosomal Protein Genes (RPGs). RSC occupancies in WT cells at the bottom 10%-Pol II occupied genes (Pol II-B10) are shown for comparison. D-E) Metagene profiles showing the normalized RSC occupancies for WT, *sth1ΔBrD, gcn5Δ,* HS-WT, *esa1* and *esa1/gcn5Δ* at RPGs (D) and at the top 20% Pol II-occupied genes excluding RPGs (E). Profiles show 500 bp upstream and 1000 bp downstream of the TSS. F) Scatterplot comparing the changes in RSC occupancies (mutant – WT; ΔRSC) in *gcn5Δ* at ORFs and promoters with WT Pol II occupancies in the ORFs of the Pol II-T20 genes (N=1092).

Next, we examined RSC occupancies at ribosomal protein genes (RPGs). RSC occupancies were higher in the ORFs of RPGs compared to the ORFs of the bottom 10% of transcribed genes (Pol II-B10) (Fig. 6C; compare Pol II-B10 with WT). The median RSC occupancies in WT (median = 2.19) and *sth1ΔBrD* (median = 2.18) were indistinguishable from each other, suggesting that the BrD is not necessary to recruit or to retain RSC over the ORFs of RPGs (Fig. 6C and 6D; left-hand panel). In contrast, the median RSC occupancy was significantly reduced in *gcn5*Δ (median = 1.36) compared to WT cells (Fig. 6C and 6D). *esa1* also reduced RSC occupancies, although less effectively than the *gcn5*Δ cells. Moreover, *esa1/gcn5Δ* showed a greater reduction in RSC than *esa1*, suggesting that the two HATs independently contribute to RSC recruitment to RPGs (Fig. 6C and 6D). It is worth noting that despite significant reductions in RSC ORF occupancies in the HAT mutants, the occupancies were still higher than that seen for the Pol II-B10 genes (Fig. 6C; compare B10 with WT and mutants). The residual binding of RSC in HAT mutants suggests the existence of additional mechanisms for recruiting RSC to the ORFs of transcribed genes. For example, RSC could bind to ORF nucleosomes by recognizing partially-unwrapped nucleosomes that are abundant in transcribed ORFs (19). In support of this idea, we have shown that RSC-bound nucleosomes in ORFs are highly accessible and resemble the RSC-bound fragile nucleosomes found in many NDRs (19,25,29). In summary, the data show that Gcn5 and Esa1 are required for WT-levels of RSC in the ORFs of RPGs.

Next, we looked at the changes in RSC enrichment in the top two deciles of transcribed genes excluding RPGs (N=966). Similar to that seen in RPGs, RSC occupancies were lowest in *gcn5*Δ (Fig. 6E; left-hand panel). The effects of *esa1* on RSC occupancies were modest compared to the *gcn5*Δ mutant for these genes, but stronger than the effect seen in the heat-shocked WT cells (Figure 6E; compare blue and gray traces in the right-hand panel). These results reinforce a greater role for Gcn5 in RSC enrichment in highly transcribed ORFs and also implicate Esa1 in this process. Surprisingly, the double mutant *esa1/gcn5Δ* did not show reduction, but rather RSC occupancies appeared to increase away from the TSS towards the 3’ end (Fig. 6E; compare solid blue and dotted black traces). This increase is inconsistent with the reduction seen in the single HAT mutants. It is plausible that the presence of both hypoacetylated or unacetylated H3 and H4 tails in the double mutant significantly stabilize RSC-nucleosome interactions, masking the actual RSC occupancy defects caused in the *esa1/gcn5Δ* mutant.

More recently, expression analyses performed after depleting the SAGA- or TFIID-specific subunits identified genes that showed reduction upon depleting TFIID, but were more sensitive to the depletion of SAGA (53) . These genes were highly expressed and were classified as “coactivator redundant (CR)” genes. We found that out of 655 CR genes, 413 genes were among the top 20% transcribed genes. The CR genes also showed a greater dependence on Gcn5 for recruiting RSC than on Esa1 (Figure S3D).

Altogether, our data implicate HATs, especially Gcn5, in maintaining high levels of RSC over transcribed ORFs. Considering the preponderance of BrD-containing subunits in the RSC complex, our data suggest that Gcn5 and Esa1 might help recruit RSC to transcribed ORFs via acetylation of histone N-terminal tails.

### The increase in RSC occupancy in promoters is independent of the loss of RSC from their ORFs in the HAT mutants

Having seen increased RSC occupancies in promoters and decreased occupancies in ORFs in the HAT mutants, we considered the possibility that RSC migrates to transcribed ORFs after being recruited to promoters in a HAT-dependent manner. Since *gcn5*Δ showed the maximal reductions in ORF RSC occupancies (Fig. 6A and 6D), we compared the reductions in the RSC occupancies in ORFs to the increases in promoters in *gcn5*Δ for the top 20% transcribed genes. In the *gcn5*Δ mutant, RSC reduction in ORFs correlated with Pol II occupancies at Pol II-T20 (Fig. 6F). In contrast, the increased RSC occupancies in the promoters correlated poorly with Pol II occupancy (Pearson R = -0.06) and with the loss of RSC from ORFs (Pearson R = 0.11) (Fig. S3E). The changes in the RSC occupancies in promoters and ORFs were also poorly correlated in other mutants (Fig. S3E). Thus, our analyses suggest that the failure of RSC to migrate to the ORFs of transcribed genes is less likely to be the major contributor to the increase in RSC seen in promoters.

### RSC recruitment to transcribed ORFs depends on histone H3 and H4 tails

The acetylation of histone tails by Gcn5 and Esa1 and the subsequent recognition of the acetylated tails by RSC bromodomains (Fig. 1A) could explain why RSC recruitment to coding sequences is reduced in cells deficient for Gcn5 and Esa1 functions. To test this, we analyzed RSC occupancies in ORFs of the Pol II-T20 genes in *H3Δ1-28* and *H4Δ1-16* mutants. The mutants showed significantly reduced RSC occupancies in ORFs of Pol II-T20 genes, RPGs, and Pol II-T20 genes excluding RPGs (Fig. 7A-C). Interestingly, the RSC occupancies were reduced to a greater extent in *H4Δ1-16* than in *esa1* (compare Fig. 7B and 7C with 6D and 6E; right-hand panels). This result raised a strong possibility that the stimulatory effect of hypoacetylated / unacetylated tails in enhancing RSC-nucleosome interactions might be masking RSC occupancy defects in *esa1* (Fig. 5). The RSC occupancy reductions were also less pronounced in the *H3Δ1-28* mutant than the *gcn5*Δ mutant. It is possible that acetylation of residues beside those in the H3 1-28 tail may also contribute to RSC recruitment. For example, H3K36 is also shown to be acetylated by Gcn5 in *S. cerevisiae* (54, 55) and could contribute in recruiting RSC. Taken together, our data strongly implies that acetylation of H3 and H4 tails promote RSC recruitment (or retention) to the gene bodies of highly transcribed genes.

**Figure 7:**
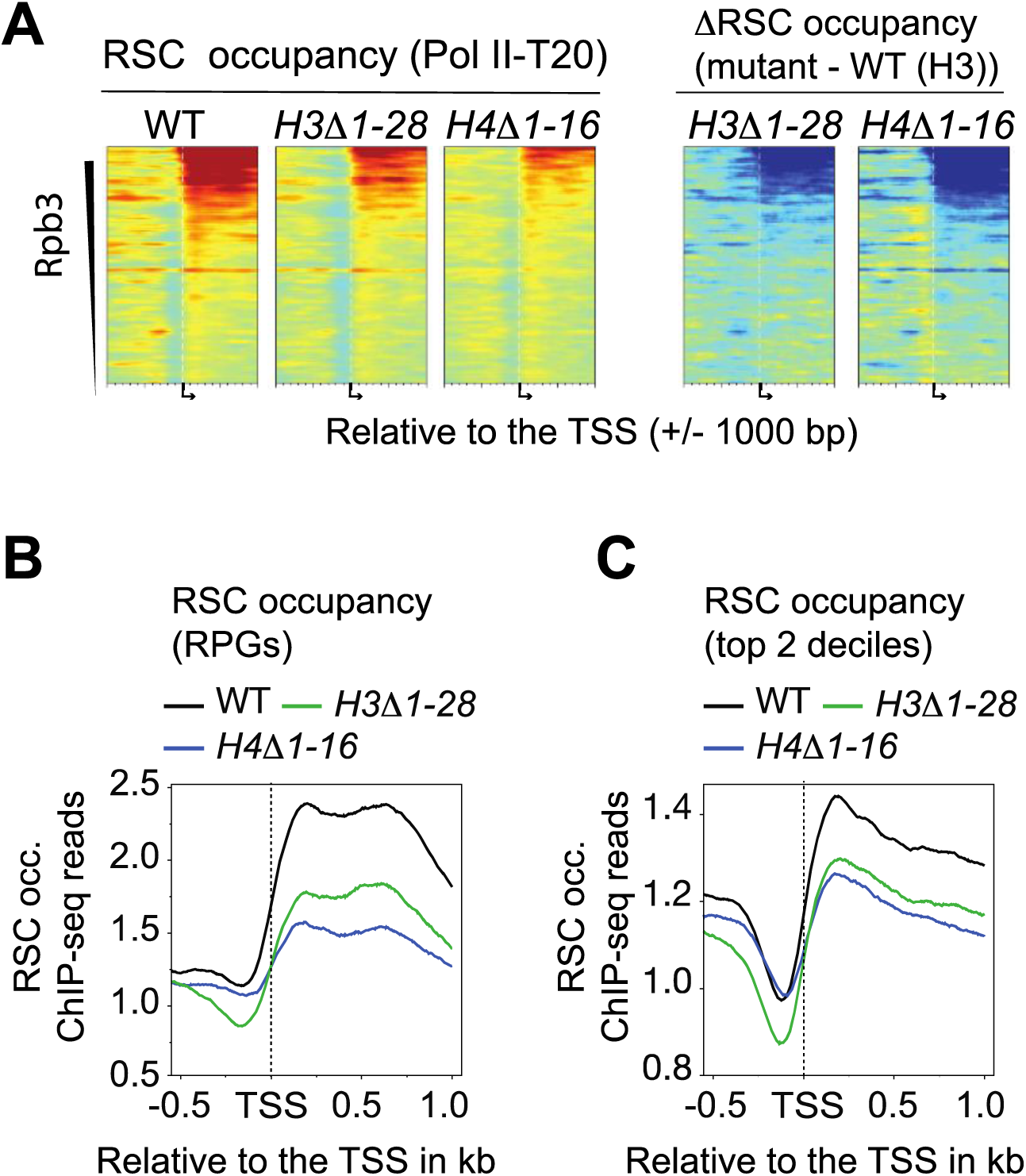
RSC recruitment to transcribed ORFs depends on H3 and H4 tails. A) Heatmaps showing RSC occupancies (RSC occ.; the three left-hand panels) and changes in RSC occupancies (mutant – WT; ΔRSC; two right-hand panels) at the top 20% Pol II-occupied genes (Pol II-T20). The heatmaps show +/- 1000 bp of the transcription start sites (TSS) for WT, *H3Δ1-28* and *H4Δ1-16* cells. Genes were sorted by decreasing average Pol II (Rpb3) occupancies in their ORFs. B-C) RSC occupancy profiles for the ribosomal protein genes (RPGs) (B) and Pol II-T20 genes, excluding RPGs (C) are shown.

### Esa1 and Gcn5 promote Pol II occupancies

Having observed that the loss of Gcn5 and Esa1 increase histone and RSC occupancies in NDRs, we next asked whether these increases affect transcription. To this end, we performed Pol II ChIP-seq using anti-Rpb3 antibodies. We observed high correlations between average Pol II occupancies in the ORFs (Pol II occupancy hereafter) of WT and HS-WT cells (R^2^=0.95; Pearson R = 0.97), indicating that Pol II occupancies were not significantly changed upon heat shock. We grouped Pol II occupancies in WT cells into deciles (D1 genes contain the most Pol II, D10 the least) and analyzed the changes in each mutant for each decile. This analysis revealed that *sth1ΔBrD* had minimal effect on reducing Pol II occupancies, except for a small reduction seen for decile 2 (D2) (Fig. 8A). In contrast, Pol II occupancies decreased significantly for genes in the first three deciles in *gcn5Δ*. The most reduction was seen for D1, which also showed the highest increase in H3 occupancies in NDRs and +1_Nuc, suggesting that the increased histone occupancies may reduce transcription of these genes. Surprisingly, Pol II occupancies appear to increase for deciles D5 to D10. A similar increase was also seen in the *sth1ΔBrD* mutant for deciles D6 to D10. Since Gcn5 promotes RSC recruitment to transcribed ORFs, the increase in Pol II occupancies might represent an elongation defect due to inefficient chromatin remodeling in ORFs. Nonetheless, the Pol II occupancies for the most highly expressed genes are reduced in cells lacking Gcn5, consistent with earlier studies (11, 56).

**Figure 8:**
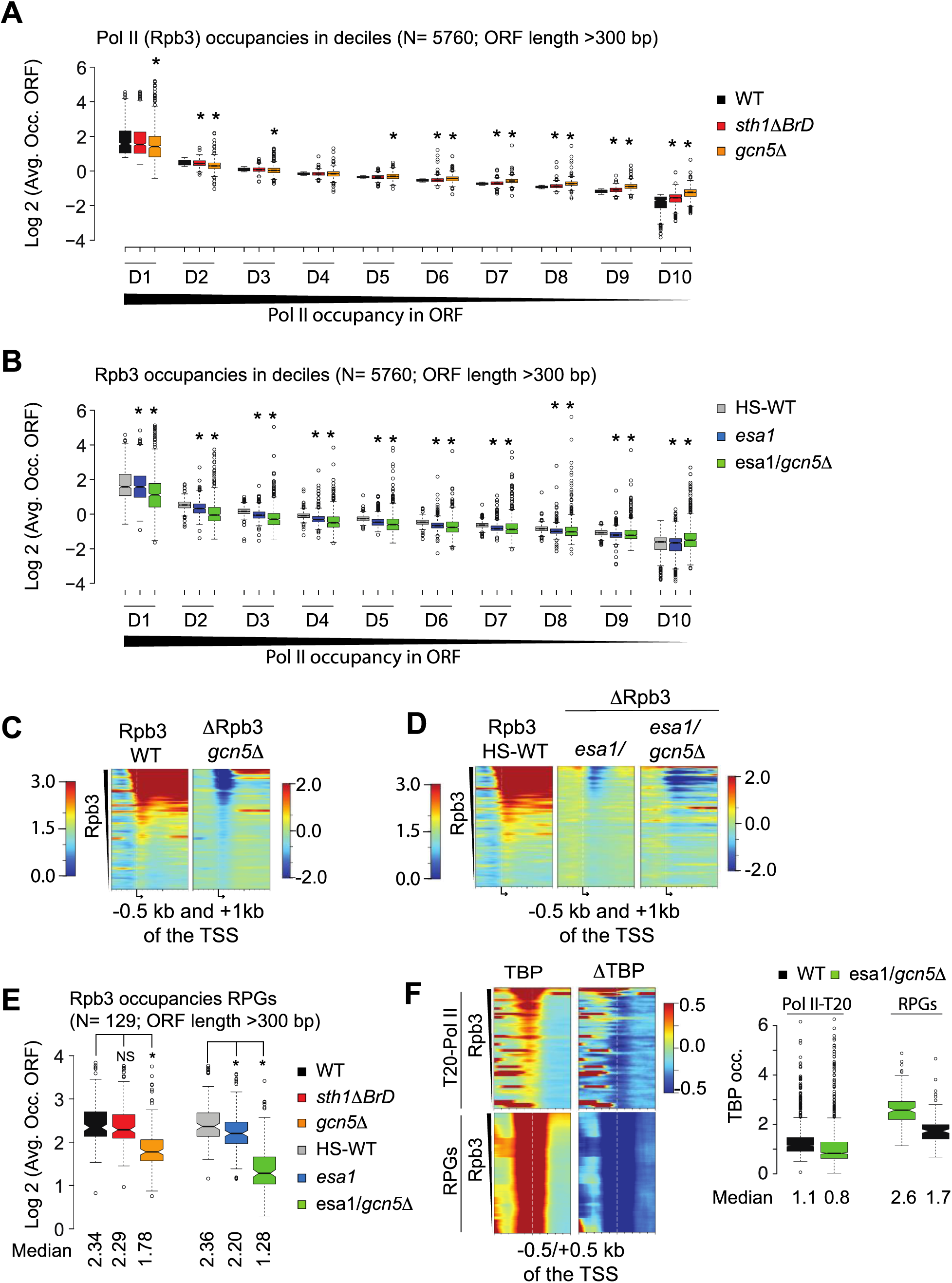
HAT mutants reduce Pol II and RBP occupancies. A-B) Boxplots depicting the average Pol II (Rpb3) occupancies (Log2) in WT and in *sth1ΔBrD* and *gcn5Δ* mutants (A), as well as in HS-WT, *esa1* and *esa1/gcn5Δ* (B). The genes were sorted based on the average Pol II occupancies in their ORFs, separated into deciles (D; N=576 per decile) such that the highest Pol II-occupied genes are in decile D1 and lowest in D10. Asterisk (*) represents significant changes (p-value < 0.05) that were determined using the Welch t-test. C-D) Heatmaps showing Rpb3 occupancies (Rpb3) and the changes in Rpb3 occupancies (ΔRpb3) at the top 20% Pol II-occupied genes (excluding RPGs; D1+D2) in WT (Rpb3) and for *gcn5Δ* (ΔRpb3) (C), and for HS-WT (Rpb3) and ΔRpb3 for *esa1* and *esa1/gcn5Δ* (D). The scales for Rpb3 are shown on the left-hand side of the heatmaps and ΔRpb3 on the right-hand side. The heatmaps show -500 bp and + 1000 bp of the TSS. E) Boxplot depicting the average Pol II (Rpb3) occupancies (Log2) at RPGs in WT, *sth1ΔBrD, gcn5Δ*, HS-WT, *esa1*, and *esa1/gcn5Δ*. The occupancies in the mutants were compared to the WT or HS-WT, and asterisks represent p-value < 0.05, calculated using the Welch t-test. NS = not significant. F) Heatmaps showing TBP occupancies (left-hand panels) and ΔTBP (right-hand panels) for the top 20% Pol II-occupied genes (D1+D2) for WT and *esa1/gcn5Δ*. The boxplots show TBP occupancy at the top two deciles of Pol II-occupied genes and for RPGs.

In contrast to *gcn5Δ*, the median Pol II occupancies for D1 decreased only slightly in *esa1* (Fig. 8B). Strikingly, however, deciles D2-D9 showed significantly reduced Pol II occupancies, suggesting that Esa1 is required for the transcription of the majority of genes. Furthermore, the double mutant *esa1/gcn5Δ* showed remarkable reductions in Pol II occupancies for deciles D1-D9. The exception was decile D10, which showed an increase. Together, these results show that Gcn5 and Esa1 are important for robust transcription of all genes, and that Gcn5 is possibly more required for transcription of highly expressed genes. It is possible that the Pol II occupancy defects in *esa1* are suppressed by Gcn5, especially considering that *esa1/gcn5Δ* double mutant showed significantly more reduction at D1 genes than the single mutants.

The heatmaps showing changes in Pol II occupancies (ΔRpb3) for the Pol II-T20 genes (D1 and D2) revealed non-uniform changes in Pol II (Fig. 8C and 8D). In *gcn5Δ*, the biggest reduction in Pol II was seen at the TSS, likely reflecting a preinitiation complex assembly defect (56). However, the genes also show a slight accumulation of Pol II in their ORFs (Fig. 8C; right-hand panel). We have shown previously that RSC inactivation also results in higher Pol II occupancies in ORFs even at genes that show lower TBP occupancy in the RSC mutant (19). *esa1* also showed greater reductions in Pol II occupancies near the TSS than in the ORFs (Fig. 8D; middle panel), and the double mutant *esa1/gcn5Δ* showed reductions throughout the ORFs (Fig. 8D, right-hand panel). The additive effects of HATs in stimulating transcription were evident at RPGs, where the double mutants showed significantly greater Pol II reductions than those seen in the single HAT mutants (Fig. 8D and 8E).

To determine whether the increase in H3 occupancies in *esa1/gcn5Δ* reduced PIC assembly, we performed TBP ChIP-seq. The TBP binding defect was significant in RPGs and in the top two deciles excluding RPGs (Fig. 8F). The reduction in TBP binding is consistent with the significant increase in H3 occupancies seen in NDRs for the highly transcribed genes.

In summary, our study revealed that HATs are important for evicting histones from the NDRs of a majority of genes. The ineffective clearance of histones from NDRs and +1 nucleosomes impede TBP recruitment and dampen Pol II occupancies. In addition, HATs are critical for RSC recruitment to transcribed ORFs in an H3 and H4 tail-dependent manner. Finally, we also uncovered a possible role for unacetylated/hypoacetylated H3 and H4 tails in the recruitment or retention of RSC to fragile nucleosomes.

## DISCUSSION

This study examined the importance of HATs Gcn5 and Esa1 in regulating occupancy of the essential chromatin remodeler RSC. The RSC ChIP-seq data revealed contrasting effects of the HATs in regulating RSC occupancies in promoters and ORFs. Deleting the H3 HAT Gcn5, inactivating the H4 HAT Esa1, or deleting either the H3 or H4 tails severely reduced RSC occupancies from highly transcribed ORFs. These results demonstrate that acetylation of histone tails by HATs is important for recruiting or retaining RSC in transcribed sequences. In contrast to the effects observed in ORFs, inactivation of HATs resulted in substantial RSC accumulation in NDRs, suggesting that histone acetylation precludes binding of RSC in NDRs. However, no accumulation is seen in the histone tail mutants. These latter observations suggest that hypoacetylated or unacetylated tails enhance retention of RSC in NDRs and suggest that acetylation of histone tails might aid in RSC disengagement. H3 ChIP-seq revealed that histones accumulate in NDRs when cells lack Gcn5 or Esa1, suggesting a global role for HATs in clearing histones from NDRs. The altered distribution of RSC and increased histones in NDRs likely contribute to the transcription defects seen in the HAT mutants.

### Hypoacetylated / unacetylated histone tails promote RSC binding to nucleosomes in NDRs

Our data show that RSC occupancies increase in promoters and NDRs in cells lacking HATs Esa1 and Gcn5. This increase was unexpected because RSC binds to acetylated nucleosomes *in vitro* (38, 43). The loss of acetylation in the HAT mutants should have decreased, not increased, RSC occupancies in NDRs. Along with the RSC increases, we also observed higher H3 occupancies in NDRs of the HAT mutants. These findings suggest that RSC accumulates on nucleosomes within NDRs. Not all NDRs are free of histones, since approximately two thousand genes have been shown to harbor fragile nucleosomes in their NDRs, many of which are RSC-bound (25,29,35). Thus, these increases might suggest that 1) fragile nucleosomes are stabilized (less DNA accessible) in the *esa1* and *gcn5*Δ mutants, resulting in higher H3 ChIP-signals, and 2) RSC is bound more efficiently to fragile nucleosomes than stable nucleosomes. We think that both increased nucleosome occupancies and increased stabilization contribute to the higher RSC occupancies found in NDRs. We find a similar increase in RSC and histones at stable nucleosome promoters in the HAT mutants, suggesting that we see more RSC because there are more histones (nucleosomes). However, at FNs, we found higher RSC occupancies and lower H3 occupancies compared to SNs, suggesting that these partially unwrapped nucleosomes retain RSC better than canonical or stable nucleosomes. One of the aspects in which the nucleosomes differ in WT and the HAT mutants is that the mutants have hypoacetylated/unacetylated tails. If hypoacetylated tails stabilize RSC occupancy, then tail-less nucleosomes should not be able to retain RSC in NDRs. Indeed, RSC did not accumulate at NDRs in cells lacking either H3 or H4 tails. Thus, our data suggest that hypoacetylated or unacetylated tails promote RSC binding to nucleosomes, this binding, however, is weakened by the acetylation of histone tails. This was surprising, given that RSC bromodomains should enhance RSC binding to nucleosomes.

Recent Cryo-EM RSC-nucleosomes structures reveal that RSC makes multiple interactions with nucleosomes involving many subunits, including Sth1 (44–46). Interestingly, one of the structures showed that the (unacetylated) H4 tail binds to the interface of the ATPase and SnAC domains of Sth1 (46). As such, the acetylation of H4 might weaken the interaction of the H4 tail with RSC and could help in RSC dissociation. The fact that RSC occupancies increase in NDRs along with histones in the HAT mutants provides *in vivo* evidence for this. Among all NDRs, RSC appears to disproportionately increase in NDRs containing fragile nucleosomes in the HAT mutants. We speculate that the partially-unwrapped fragile nucleosomes might allow better association of the H4 tail with the Sth1 interface than more compactly wrapped (stable) nucleosomes at -1 positions. We speculate that the hypo/unacetylated tails of FNs attract more RSC to facilitate the eviction of promoter nucleosomes and allow these genes to achieve higher transcription levels. A recent study showed that RSC binds to NDRs in quiescent cells and that this binding was deemed critical for hyper-transcription (57). We speculate that the ability of FNs to recruit RSC in a largely acetylation-independent manner might prime these promoters to respond to transcription-activating signals rapidly.

### HATs prevent histone accumulation in NDRs genome-wide

Chromatin remodelers are implicated in the formation and maintenance of canonical chromatin structure in which NDRs are flanked by highly positioned -1 and +1 nucleosomes (20,30,58,59). The RSC complex is particularly important in establishing NDR width genome-wide (21,23,58,60). It is widely accepted that histones are evicted from promoters during the induction of genes (27,61–63). For example, studies have shown that deletion of *GCN5* increases histone levels at the promoters of genes that are induced by sulfometuron methyl (SM) (10, 11), a chemical that inhibits isoleucine/valine biosynthesis and causes amino starvation stress. The highest increase in histones was observed in the NDRs of SM-induced genes. Our studies show that H3 levels increased in most NDRs, including those of the least transcribed genes. The global H3 occupancy increases in NDRs in *gcn5Δ*, *esa1* and *esa1/gcn5Δ* suggests that both HATs and RSC play a role in keeping NDRs depleted of nucleosomes. Interestingly, *sth1ΔBrD* also showed increased H3 occupancies in NDRs, and these increases were very similar to that seen in *gcn5Δ* cells for the bottom 75% transcribed genes (Fig. S2B). It makes sense considering that acetylated H3K14, a Gcn5 target, is recognized by the Sth1 bromodomain (42,64,65).

The fact that HAT mutants increase both RSC and histone occupancies makes it tempting to speculate that although hypoacetylated histones in NDRs can attract more RSC, the unacetylated histones are not effectively evicted. In fact, biochemical experiments have shown that acetylation of histone tails stimulates RSC catalytic activity *in vitro* (38, 39). Thus, FNs can recruit RSC in an acetylation-independent, but tail-dependent manner and that the subsequent acetylation of tails might stimulate RSC to remove these nucleosomes more efficiently. Rapid RSC recruitment and histone eviction from promoters could explain why genes with FNs have more Pol II in their ORFs than SN genes. The RSC accumulation in NDRs seen in the HAT mutants but not in the tail mutants suggests that RSC can be recruited to nucleosomes using hypoacetylated (or unacetylated) tails and that subsequent acetylation might help RSC to evict histones (38,39,42,65).

### Acetylated histone H3 and H4 tails promote RSC occupancy in transcribed ORFs

Acetylation is thought to promote transcription by weakening histone tail-DNA interactions. However, a recent study challenged this and showed that acetylation of histone tails is promoted by transcription (12). Acetylation of histone tails is also thought to promote the binding of bromodomain-containing complexes that can relieve the nucleosomal barrier for transcription (13,15,66). Our data support this idea in that HATs are important for enriching RSC in transcribed coding sequences. Loss of Gcn5 severely dampened RSC occupancies in transcribed regions (Fig. 6A, 6C-E). In comparison, the loss of Esa1 activity evoked only a small reduction in RSC occupancy from ORFs. Notwithstanding modest reductions in RSC occupancies in *esa1*, the H4 tail mutant (*H4Δ1-16*) showed a substantial reduction of RSC from transcribed ORFs (Fig. 7), and this was also seen in the H3 tail mutant (*H3Δ1-28*), providing strong support for histone tails in mediating RSC recruitment/retention in ORFs. As such, the data strongly suggest that one of the functions of transcription-dependent acetylation of ORF nucleosomes is to recruit or maintain high levels of RSC occupancies in the transcribed sequences. Recruitment of RSC to ORFs might make nucleosomes more accessible to elongating RNA polymerase. Consistent with this, we have shown that RSC-bound nucleosomes in the ORFs are digested to a greater extent than non-RSC-bound nucleosomes (19).

### HATs promote transcription

Pol II occupancy analyses in cells lacking Gcn5 or Esa1 function reveal that these HATs have a differential effect on Pol II occupancies. *gcn5Δ* cells evoked Pol II reduction from highly transcribed genes, whereas the *esa1* mutant displayed Pol II reduction for most genes, except for the top 10% and the bottom 10% of highly transcribed genes. This suggests that Esa1 is more broadly required for Pol II occupancies than Gcn5 in facilitating transcription, consistent with previous studies (53,56,67). The double mutant *esa1/gcn5Δ* showed the greatest reductions in Pol II occupancies for ∼90% of the transcribed genes. A greater Pol II occupancy defect in the double mutant compared to the single mutants at Pol II-T10 genes (Fig. 8A-B) suggest that Gcn5 might be compensating for the loss of Esa1 function on transcription at some genes, but it is not able to do so at other genes. Additionally, the double mutant also revealed a role for Gcn5 on genes other than the most highly expressed since the double mutant showed greater reductions than those seen in *esa1* for the majority of the genes. Since both HATs are important for enriching RSC in ORFs, it is possible that the slower migration of Pol II through genic regions dampens the actual transcription defects caused by the absence of HATs. The double mutant also showed a significant reduction in TBP occupancies for RPG genes and for the top 20% of transcribed genes, suggesting that increased histone occupancies in NDRs interfere with TBP binding and possibly PIC formation.

Our data suggest that both RSC and HATs are important for evicting histones from NDRs and promoting PIC assembly and transcription. We propose a model in which HATs are important for evicting histones from NDRs, genome-wide (Fig. 9). The unacetylated tails might recruit RSC to NDRs, and subsequent acetylation of these tails by HATs Gcn5 and Esa1 leads to histone eviction from the NDR to promote PIC assembly and transcription (Fig. 9A). However, in cells lacking HATs, RSC is still recruited to NDRs, but histone eviction is reduced. These increased histone levels in NDRs lead to reduced TBP binding and Pol II recruitment, resulting in dampened transcription (Fig. 9B).

**Figure 9:**
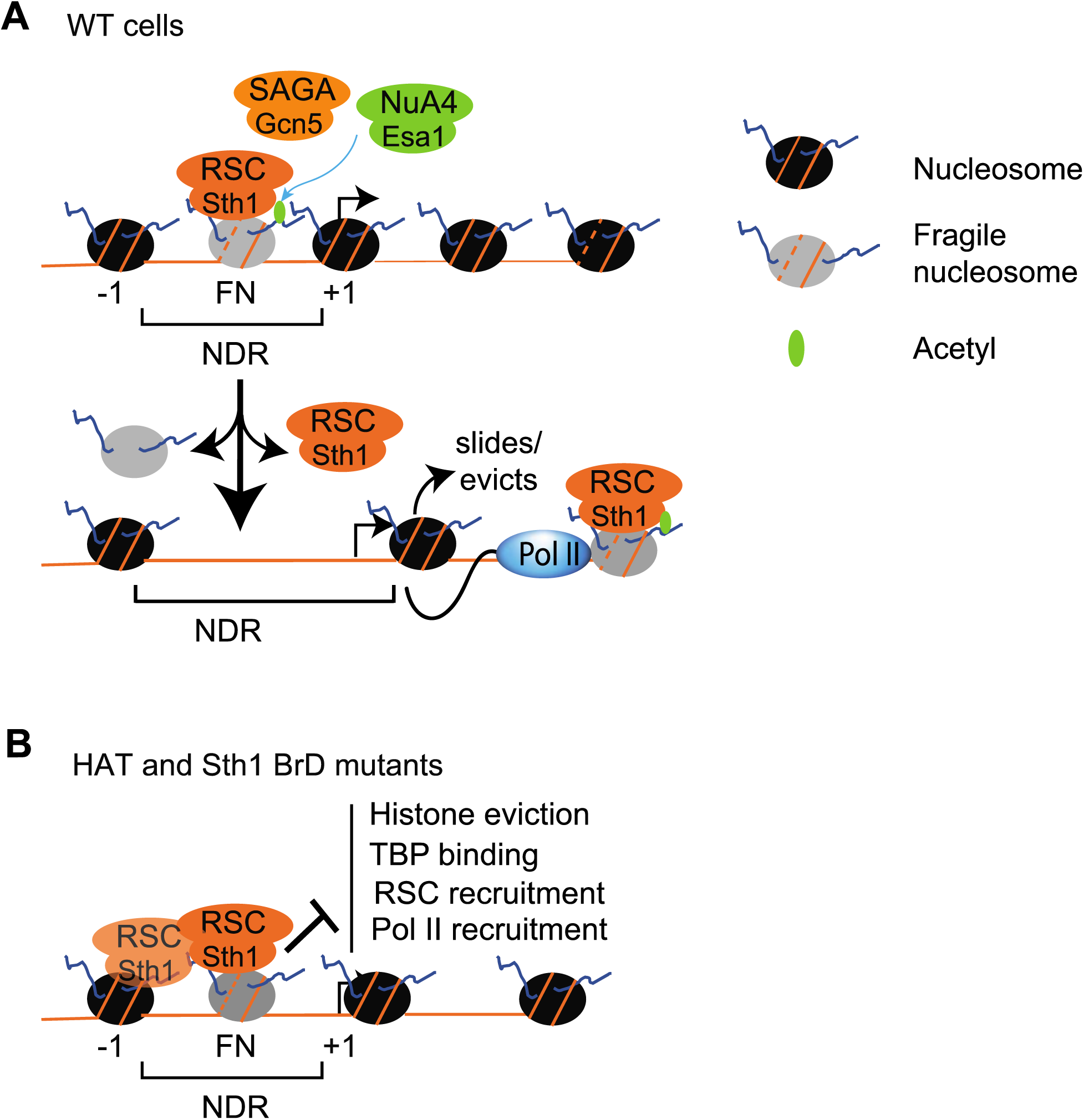
Model depicting the roles for HATs (Gcn5 and Esa1) in regulating chromatin structure and transcription. A) At a typical gene, the NDR is flanked by +1 and -1 nucleosomes. Some genes with wider NDRs also harbor partially-unwrapped nucleosomes such as fragile nucleosomes that have been shown to be RSC bound (29, 35). In WT cells, histones are evicted by RSC, which could be recruited to the NDR through transcription activators, histone acetylation by either Gcn5 or Esa1, or by recognizing specific DNA sequences. RSC can either evict the +1 nucleosome or slide it in the 3’ direction to expose the TBP binding site, leading to increased TBP and Pol II recruitment in promoters. In addition, RSC is also recruited to transcribed ORFs in a transcription dependent manner. B) Our data implicate hypoacetylated/unacetylated H3 and H4 tails in stabilizing RSC on fragile nucleosomes and also on some stable nucleosomes. The nucleosomes in NDRs are less efficiently removed when histones are hypoacetylated/unacetylated, as in the Gcn5 and Esa1 mutants, leading to reduced TBP binding and Pol II recruitment. Altogether, our data suggest that fragile nucleosomes might use hypo- or unacetylated histone tails to recruit RSC to FNs, and subsequent acetylation of these tails by HATs stimulates histone eviction. Thus, HATs, in coordination with RSC, help maintain NDRs genome-wide and also helps genes to achieve high levels of transcription by promoting TBP recruitment and possibly by aiding in Pol II elongation by promoting RSC recruitment in ORFs.

## DATA AVAILABILITY STATEMENT

The accession numbers for the raw and analyzed data reported in this paper are GSE206405. The strains are available upon request.

## FUNDING

This work was supported by the National Institutes of Health R01GM095514 and R15GM126449 to CKG, and from Center for Biomedical Sciences, Oakland University.

## ACKNOWLEDGEMENT

We are grateful to Dr. Joseph Resse (Penn State University) for providing anti-TBP antibodies. We thank Dr. Alan Hinnebusch for providing *esa1/gcn5Δ* mutant strain, and for providing useful comments on the manuscript.

## Notes

### Competing Interest Statement

The authors have declared no competing interest.

