## Supplemental figures for "The role of histone acetyltransferases Gcn5 and Esa1 in recruiting the RSC complex and maintaining nucleosome-depleted regions genome-wide in *Saccharomyces cerevisiae*"

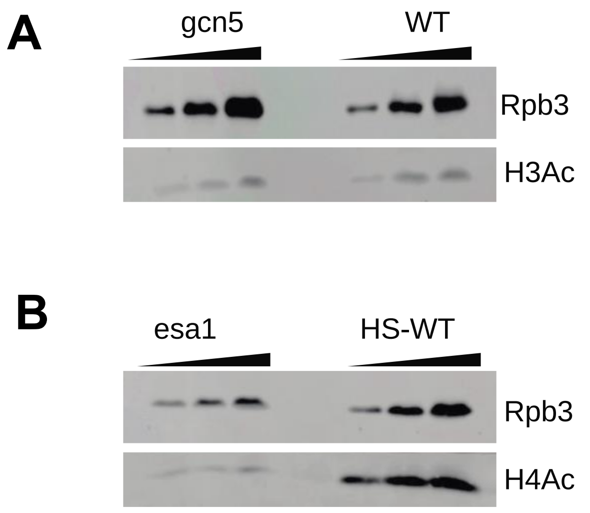


Figure S1

**Figure S1: Reduced H3 and H4 acetylation in *gcn5*Δ and *esa1ts* mutant.**

A-B) Western blots showing the levels of H3 acetylation (H3Ac) in WT and in *gcn5Δ* (A) cells, and H4 acetylation (H4Ac) in HS-WT cells and in the *esa1* mutant. The different lanes for the indicated strain show different amount of whole cell extract loading. Pol II (Rpb3) was used as the loading control.

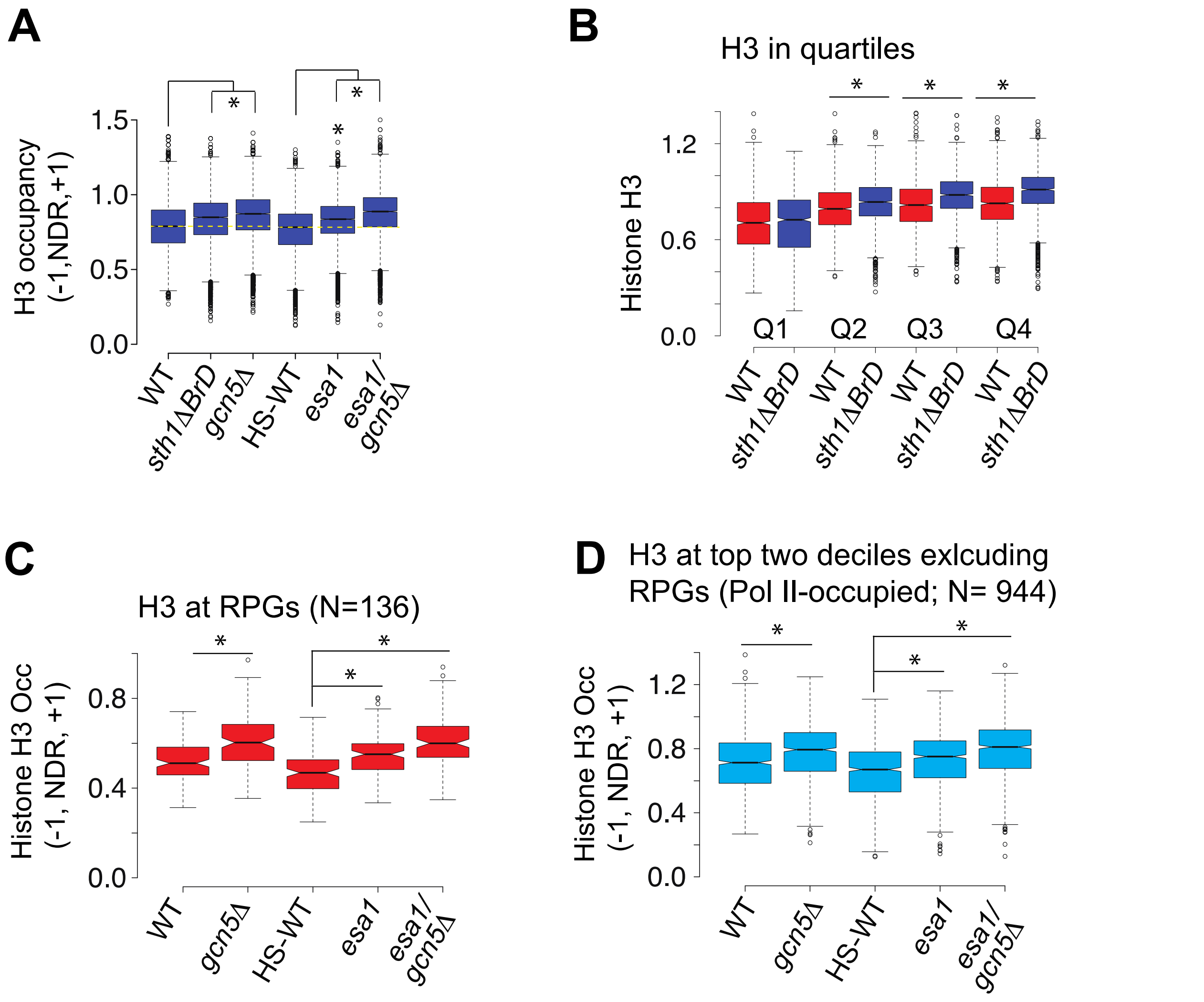


Figure S2

**Figure S2: H3 occupancies increase in the promoters of HAT and Sth1 BrD mutants.**

A) Boxplot showing the normalized H3 occupancies at promoters (-1_Nuc, NDR, and +1_Nuc positions) in WT cells, HS-WT cells, *sth1ΔBrD*, *gcn5Δ*, *esa1* and *esa1/gcn5Δ* cells. The dotted yellow line represents the median of WT and HS-WT H3 occupancies.

B) Boxplot showing the H3 occupancies at promoters (-1_Nuc, NDR, and +1_Nuc positions) in quartiles for WT and *sth1ΔBrD*. H3 significantly increases in *sth1ΔBrD* mutant for quartiles Q2 to Q4. The quartiles are based on the average Pol II occupancies in gene ORFs, where genes in Q1 harbor the most Pol II and genes in Q4 harbor the least.

C-D) H3 occupancy in promoters of RPGs (C) and of the top two deciles of Pol II-occupied genes excluding RPGs (D).

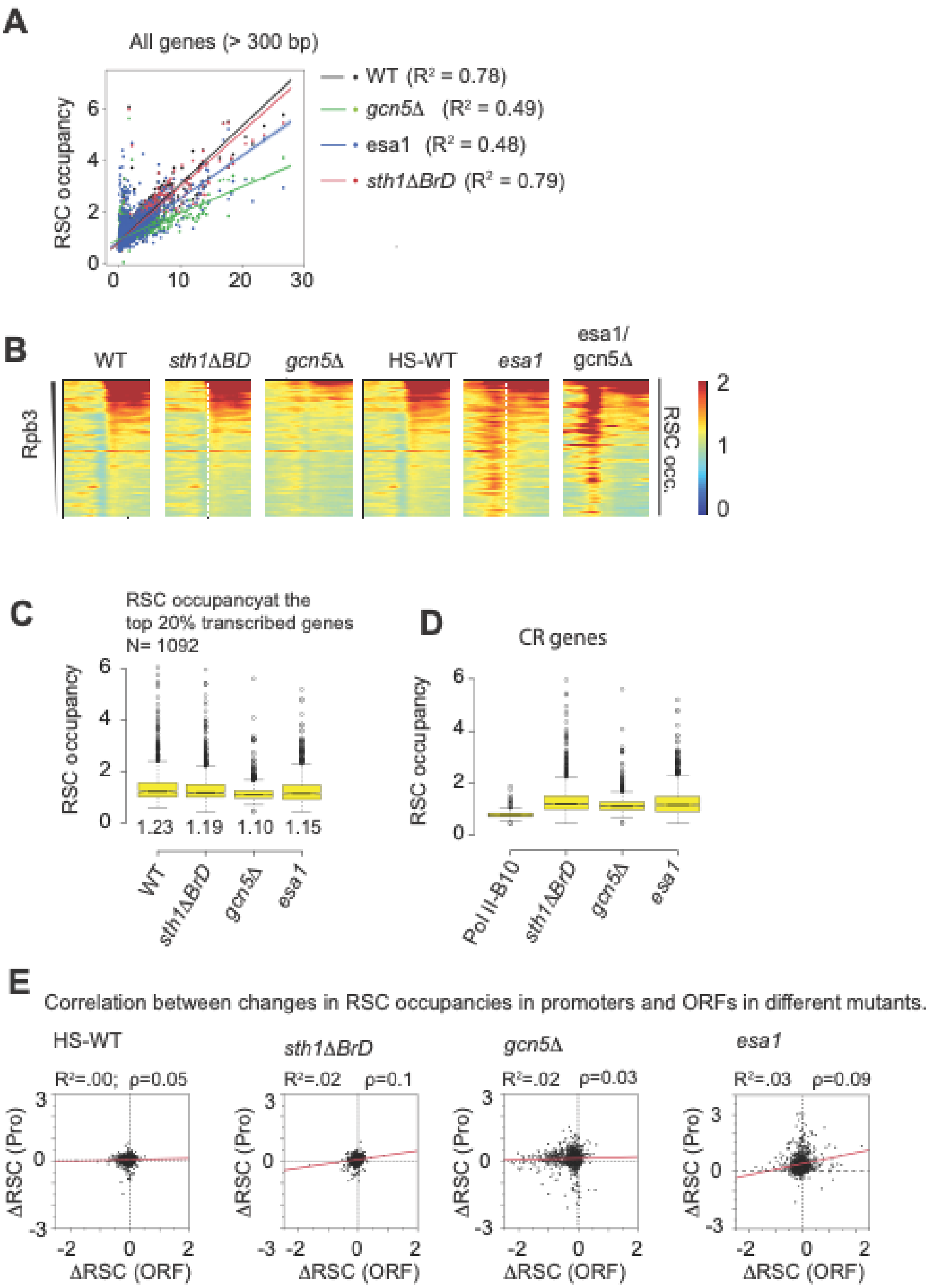

A
All genes (> 300 bp)
RSC occupancy
WT (R² = 0.78)
gcn5Δ (R² = 0.49)
esa1 (R² = 0.48)
sth1ΔBrD (R² = 0.79)
B
WT
sth1ΔBD
gcn5Δ
HS-WT
esa1
esa1/gcn5Δ
Rpb3
RSC occ.
C
RSC occupancyat the top 20% transcribed genes N= 1092
RSC occupancy
1.23 1.19 1.10 1.15
WT
sth1ΔBrD
gcn5Δ
esa1
D
CR genes
RSC occupancy
Pol II-B10
sth1ΔBrD
gcn5Δ
esa1
E
Correlation between changes in RSC occupancies in promoters and ORFs in different mutants.
HS-WT
R²=.00; ρ=0.05
sth1ΔBrD
R²=.02 ρ=0.1
gcn5Δ
R²=.02 ρ=0.03
esa1
R²=.03 ρ=0.09
ΔRSC (Pro)
ΔRSC (ORF)


Figure S3

**Figure S3: HATs promote RSC recruitment to transcribed ORFs.**

A) Scatter plots showing correlation between RSC occupancies and Pol II occupancies in WT, *sth1ΔBrD*, *gcn5Δ* and *esa1* cells.

B) Heatmaps showing RSC occupancies at top 20% Pol II-occupied genes (Pol II-T20) for WT, *gcn5Δ*, *sth1ΔBrD*, HS-WT, *esa1* and *esa1/gcn5Δ*. Genes were sorted by decreasing average Pol II (Rpb3) occupancies in their ORFs. The heatmaps show +/- 1000 bp of the transcription start sites (TSS).

C-D) Boxplots showing RSC occupancies for WT, *sth1ΔBrD, gcn5Δ, and esa1* at Pol II-T20 genes (C) and for coactivator regulated (CR) genes (D). RSC occupancies in WT cells at the bottom 10%-Pol II occupied genes (Pol II-B10) are shown for comparison.

E) Scatter plots showing that RSC occupancy changes in promoters (ΔRSC (Pro)) poorly correlates with changes in ORF occupancies (ΔRSC (ORF)). The $R^2$ and Spearman correlation (ρ) are shown for each comparison.

Table S1

Title: List of strain used in the study

| Name | Parent | Genotype | Reference |
|---|---|---|---|
| F729 | BY4741 | *MATa his3Δ1 leu2Δ0 met15Δ0 ura3Δ0* | Research Genetics |
| CGY 2 | BY4741 | *MATa his3Δ1 leu2Δ0 met15Δ0 ura3Δ0 STH1-myc13::HIS3* | This study |
| DG 265 | 7285 | *MATa his3Δ leu2Δ met15Δ ura3Δ gcn5Δ::kanMX4 STH1-myc13::HIS3* | (1) |
| DG 259 | DG150 | *MATa his3Δ leu2Δ met15Δ ura3Δ esa1L254P STH1-myc13::HIS3* | (1) |
| DG 271 | DG154 | *MATa his3Δ leu2Δ met15Δ ura3Δ gcn5Δ::kanMX4 esa1L254P STH1-myc13::HIS3* | (1) |
| EBY 1 | BY4741 | *MATa his3Δ1 leu2Δ0 met15Δ0 ura3Δ0 STH1ΔBrD(1270-1359)-myc13::HIS3* | This study |
| JDY 86 | S288C | *MATa his3Δ200 leu2Δ0 lys2Δ0 trp1Δ63 ura3Δ0 met15Δ0 hht1-hhf1::NatMX4 can1::MFA1pr-HIS3 [pJP11, HHT1-HHF1 CEN-LYS2]* | (2) |
| SJY 1 | JDY 86 | *MATa his3Δ200 leu2Δ0 lys2Δ0 trp1Δ63 ura3Δ0 met15Δ0 can1::MFA1pr-HIS3 hht1-hhf1::NatMX4 hht2-hhf2::H3-URA3 STH1-myc13::TRP1* | This study |
| SJY 2 | JDY 86 | *MATa his3Δ200 leu2Δ0 lys2Δ0 trp1Δ63 ura3Δ0 met15Δ0 can1::MFA1pr-HIS3 hht1-hhf1::NatMX4 hht2-hhf2::[H3Δ1-28]-URA3 STH1-myc13::TRP1* | This study |
| EBY 11 | JDY 86 | *MATa his3Δ200 leu2Δ0 lys2Δ0 trp1Δ63 ura3Δ0 met15Δ0 can1::MFA1pr-HIS3 hht1-hhf1::NatMX4 hht2-hhf2::[H4Δ1-16]-URA3 STH1-myc13::TRP1* | This study |
